# LSD-induced increase of Ising temperature and algorithmic complexity of brain dynamics

**DOI:** 10.1101/2022.08.27.505518

**Authors:** Giulio Ruffini, Giada Damiani, Diego Lozano-Soldevilla, Nikolas Deco, Fernando E. Rosas, Narsis A. Kiani, Adrián Ponce-Alvarez, Morten L. Kringelbach, Robin Carhart-Harris, Gustavo Deco

**Affiliations:** Neuroelectrics Barcelona, Av. Tibidabo 47b, 08035 Barcelona, Spain; Starlab Barcelona, Av. Tibidabo 47b, 08035 Barcelona, Spain; Haskins Laboratories, 300 George Street, New Haven, CT, 06511, USA; Centre For Psychedelic Research (Department of Medicine), Imperial College London, London, UK; Data Science Institute, Imperial College London, London, UK; Centre for Complexity Science, Imperial College London, London, UK; Department of Informatics, University of Sussex, Brighton, UK; Algorithmic Dynamics Lab, Center of Molecular Medicine, Karolinksa Institutet, Stockholm, Sweden; Oncology and pathology Department, Karolinksa Institutet, Stockholm, Sweden; Center for Brain and Cognition, Computational Neuroscience Group, Department of Information and Communication Technologies, Universitat Pompeu Fabra, Roc Boronat 138, Barcelona, 08018, Spain; Centre for Eudaimonia and Human Flourishing, University of Oxford, Oxford, UK; Department of Psychiatry, University of Oxford, Oxford, UK; Center for Music in the Brain, Department of Clinical Medicine, Aarhus University, Aarhus, Denmark; Institucio Catalana de Recerca i Estudis Avançats (ICREA), Universitat Pompeu Fabra, Computational Neuroscience, Barcelona, Spain; Institució Catalana de la Recerca i Estudis Avancats (ICREA), Passeig Lluís Companys 23, Barcelona, 08010, Spain; Department of Neuropsychology, Max Planck Institute for Human Cognitive and Brain Sciences, 04103 Leipzig, Germany; School of Psychological Sciences, Monash University, Melbourne, Clayton, VIC 3800, Australia

## Abstract

A topic of growing interest in computational neuroscience is the discovery of fundamental principles underlying global dynamics and the self-organization of the brain. In particular, the notion that the brain operates near criticality has gained considerable support, and recent work has shown that the dynamics of different brain states may be modeled by pairwise maximum entropy Ising models at various distances from a phase transition, i.e., from criticality. Here we aim to characterize two brain states (psychedelics-induced and placebo) as captured by functional magnetic resonance imaging (fMRI), with features derived from the Ising spin model formalism (system temperature, critical point, susceptibility) and from algorithmic complexity. We hypothesized, along the lines of the entropic brain hypothesis, that psychedelics drive brain dynamics into a more disordered state at a higher Ising temperature and increased complexity. We analyze resting state blood-oxygen-level-dependent (BOLD) fMRI data collected in an earlier study from fifteen subjects in a control condition (placebo) and during ingestion of lysergic acid diethylamide (LSD). Working with the automated anatomical labeling (AAL) brain parcellation, we first create “archetype” Ising models representative of the entire dataset (global) and of the data in each condition. Remarkably, we find that such archetypes exhibit a strong correlation with an average structural connectome template obtained from dMRI (*r* = 0.6). We compare the archetypes from the two conditions and find that the Ising connectivity in the LSD condition is lower than the placebo one, especially at homotopic links (interhemispheric connectivity), reflecting a significant decrease of homotopic functional connectivity in the LSD condition. The global archetype is then personalized for each individual and condition by adjusting the system temperature. The resulting temperatures are all near but above the critical point of the model in the paramagnetic (disordered) phase. The individualized Ising temperatures are higher in the LSD condition than the placebo condition (*p* = 9 × 10^-5^). Next, we estimate the Lempel-Ziv-Welch (LZW) complexity of the binarized BOLD data and the synthetic data generated with the individualized model using the Metropolis algorithm for each participant and condition. The LZW complexity computed from experimental data reveals a weak statistical relationship with condition (*p* = 0.04 one-tailed Wilcoxon test) and none with Ising temperature (*r*(13) = 0.13, *p* = 0.65), presumably because of the limited length of the BOLD time series. Similarly, we explore complexity using the block decomposition method (BDM), a more advanced method for estimating algorithmic complexity. The BDM complexity of the experimental data displays a significant correlation with Ising temperature (*r*(13) = 0.56, *p* = 0.03) and a weak but significant correlation with condition (p = 0.04, one-tailed Wilcoxon test). This study suggests that the effects of LSD increase the complexity of brain dynamics by loosening interhemispheric connectivity—especially homotopic links. In agreement with earlier work using the Ising formalism with BOLD data, we find the brain state in the placebo condition is already above the critical point, with LSD resulting in a shift further away from criticality into a more disordered state.

**Author summary:** In this study, we aim to characterize two brain states (psychedelics-induced and placebo), as captured in functional magnetic resonance imaging (fMRI) data, with features derived from the Ising model formalism (system temperature, critical point, susceptibility) and from algorithmic complexity. Under the hypothesis that psychedelics drive the brain into a more disordered state, we study criticality features of brain dynamics under LSD in a within-subject study using the Ising model formalism and algorithmic complexity using Lempel-Ziv and the Block Decomposition methods. Personalized Ising models are created by first using BOLD fMRI data from all the subjects and conditions to create a single Ising “archetype” model—which we can interpret as the average model of the data at unit temperature—and then by adjusting the model temperature for each subject and condition. We find that the effects of LSD translate into increased BOLD signal complexity and Ising temperature, in agreement with earlier findings and predictions from existing theories of the effects of psychedelics, such as the relaxed beliefs under psychedelics (REBUS), the anarchic brain hypothesis [1], and the algorithmic information theory of consciousness (KT) [2, 3]. However, in contrast with some of the previously cited theories, we find that the system in the placebo condition is already in the paramagnetic phase—above the critical point—with ingestion of LSD resulting in a shift away from Ising criticality into a more disordered state. Finally, we highlight the fact that the structural connectome can be recovered to a good degree by fitting an Ising model and that the reduction of homotopic links appears to play an important role in the slide to disorder under psychedelics.

## Introduction

The complex features displayed by the emergent neural dynamics of the human brain are in many ways reminiscent of behaviors of paradigmatic systems in statistical physics [4–12]. For example, just as condensed materials can transit between ordered (e.g., solid) or disordered (e.g., gas) phases, neural networks can undergo phase transitions between highly regular and highly unsynchronized activity. Of particular interest are dynamics of systems poised close to phase transitions, where so-called critical points display a balance of order and disorder that has been conjectured to be crucial for living organisms [13]. At critical points, non-trivial collective patterns spanning all scales are observed, giving rise to a rich repertoire of short- and long-range correlations. The fluctuations in critical systems are highly structured, obeying fundamental physical principles rooted in the system’s symmetries—which are shared by many systems despite their underlying constituency, establishing ‘universality classes’ of large-scale coordinated activity [14].

Theoretical work indicates that computation and information dynamics in complex systems display special features at critical points [15–19]). Building on these insights, criticality has been suggested as a useful principle to account for the brain’s inherent complexity that is required to guide behavior in rich environments by processing various sources of information [5, 20–26]. Furthermore, it has been found that systems in nature often tune themselves into a critical state—a principle known as ‘self-organized criticality’ [27–29]. It has been proposed that neural systems may self-organize, evolutionary and developmentally, through homeostatic plasticity rules that result in the system arriving near critical states [30–32].

Building on this pioneering work, this paper aims to leverage tools from the statistical physics literature to study brain dynamics that support different states of consciousness. Here we focus on the altered state of consciousness induced by LSD, a psychedelic substance that is recently harnessing attention for its therapeutic potential for addressing mental disorders such as anxiety, depression, or substance addiction [33–37]. While our understanding of how LSD work at the neuronal and pharmacological levels is relatively well-developed [32, 38, 39], its effects at the whole-brain level are less understood. Yet, this is paramount to explain better and predict the outcomes of psychedelic interventions—such as psychedelic-assisted therapy [33, 40, 41]. It has been argued that the observed expansion of the repertoire of functional patterns elicited by hallucinogenic substances can be understood as being driven by an enhancement of brain entropy [32]. In turn, this may take place while the brain is moving closer to criticality, with this being related to the relaxation of high-level cognitive priors [1, 3], which in turn may lead to a favorable context for conducting psychotherapy [1, 42]. Studies on functional neuroimaging regarding LSD effects have shown initial evidence of the mechanistic alterations on brain dynamics at the network level, with the majority of the findings suggesting a relative weakening of usual functional configurations giving place to an increase of brain entropy, global functional integration, and more flexible brain dynamics [32, 33, 43–48].

Given the theoretical and empirical evidence suggesting that LSD induces a shift towards a broader repertoire of neural patterns enabled by more flexible dynamics, the present study aims to characterize the underlying neural mechanisms responsible for these effects via mathematical and computational modeling tools from statistical physics and the study of phase transitions. Specifically, we leverage maximum entropy models—the most parsimonious models one can build following statistical physics principles [49, 50]—to describe the coordinated activity of various brain areas as captured by BOLD signals. By building individualized computational models for LSD and placebo conditions, we aim to analyze differences in terms of the model’s temperature, a parameter that describes the system’s randomness. In particular, a low temperature amplifies distinctions between energy levels and hence a higher dissimilarity between the likelihood of occupancy of different states; conversely, a high system temperature ‘flattens’ the energy landscape and allows access to more states.

Moreover, the system’s temperature situates the state in relation to the critical boundaries defining phase transitions. Finally, given the intuitive relationship between signal diversity, algorithmic complexity, and criticality, we explore complexity metrics characterizing the two conditions in the data with the hypothesis that complexity correlates with system temperature and hence experimental condition [2, 3, 51].

## Materials and methods

### Experimental protocol and BOLD fMRI data

Our dataset consists of BOLD time series obtained under LSD effects and in a placebo condition from fifteen participants (N=15). Details of the data acquisition process can be found in detail in [33, 52], and we only provide a summary here. The study was approved by the National Research Ethics Service committee London-West London and was conducted in accordance with the revised declaration of Helsinki (2000), the International Committee on Harmonization Good Clinical Practice guidelines, and National Health Service Research Governance Framework. Imperial College London sponsored the research, which was conducted under a Home Office license for research with Schedule 1 drugs. Written informed consent was given, and data was de-personalized (anonymized) prior to analysis. A group of 15 participants (4 women; mean age, 30.5±8.0) were recruited by personal spoken communication and provided with written consent to participate in the experiment after a briefing and screening study for physical and mental health. This included an electrocardiogram (ECG), urine and blood tests, plus a psychiatric test and drug history. Participants under 21 years, with psychiatric diseases, family history of psychotic disorders, experience with psychedelics, pregnancy, problems with alcohol (>40 units per week), or any other medically significant condition that could affect the study were excluded. Participants attended twice the laboratory (once per condition), with a separation of 2 weeks. The order of these was alternated across participants without giving the information about the experimental condition used in each session. With the help of a medical doctor, a cannula was inserted into a vein in the antecubital fossa after the tests. The dose consisted of 75 mg of LSD via a 10 ml solution proportioned over two minutes with an infusion of saline afterward. Similarly, a placebo (10 ml saline) was injected intravenously over two minutes. After this, participants were suggested to lie relaxed with closed eyes inside the MRI scanner. Participants provided subjective reports of the drug effects after a 5 to 15 minutes period from the administration of the drug. The peak effects were reported to occur 60–90 minutes after the dose. Subsequently, effects were generally constant for four hours. The scanning was done approximately 70 minutes after the dose and lasted a full hour. BOLD scanning was composed of three resting state scans of seven minutes. The middle one incorporated listening to two excerpts of music from two songs by ambient artist Robert Rich. For our analysis here, we exclude the scan with music. Thus, the dataset includes four scans, each of 7 minutes (two under LSD, two in the placebo condition) for each of the fifteen subjects (for a total of sixty). The data from the two scans in each condition are combined in the following analysis.

### Parcellated data series

The fMRI BOLD data was preprocessed using FSL tools (FMRIB’s Software Library, www.fmrib.ox.ac.uk/fsl) with standard parameters and not discarding any Independent Component Analysis (ICA) components. This included corrections for head motion during the scans, which was within the normal, acceptable range for fMRI sessions. Also, FSL was used to generate parcellated (region-averaged) BOLD signals for each participant and condition, i.e., the signal was averaged over all voxels within each region defined in the automated anatomical labeling (AAL) atlas. Only the 90 AAL atlas cortical and subcortical non-cerebellar brain regions of interest (ROIs or parcels) were considered [53] (see Table SI.1 in Appendix A). For each subject and condition (LSD or placebo), we obtained a BOLD time series matrix of dimensions 90 x 217 (AAL regions x time points, where the sampling cadence was 2 s) from each session. The preprocessing pipeline of the BOLD data and the generation of the BOLD time series is described in more detail in [52]. See Supplementary Information for plots of group FC and FC of each subject.

### Data binarization

To build an Ising model from the data or carry out complexity analysis, we transformed each parcel’s BOLD fMRI data series into a binary format. This was done by using a threshold: each value of the region-averaged BOLD time series was assigned a value of +1 if its value was greater than the threshold and —1 otherwise. The threshold was set to the median of the time series in the parcel of interest. Hence the median of the thresholded time series of each voxel was zero, and the entropy of each time series was maximal. The binarization was carried out independently for each condition, session, subject, and parcel. The binarized data from the two scans in each session were concatenated.

### Data analysis of fMRI series

#### Lempel-Ziv complexity

Lempel-Ziv (LZ) refers to a class of adaptive dictionary compression algorithms that parse a string into words and use increasingly long reappearing words to construct a dictionary. Compression is achieved by pointing to the identifiers of words in the dictionary [54–56]. LZ algorithms are universally optimal—their asymptotic compression rate approaches the entropy rate of the source for any stationary ergodic source [56], 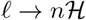. where *l* is the length of the compressed string, *n* is the length of the string and 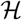 is the entropy rate.

LZ has been used across many studies to characterize brain state—from neurodegeneration [57–60], to anesthesia [61], disorders of consciousness [62, 63] to fetal consciousness [64]. The entropic brain hypothesis [44] proposes that within a critical zone, the entropy of spontaneous brain activity is associated with various dimensions of subjective experience and that psychedelics acutely increase both [1]. More recently, it has been proposed that algorithmic versions of entropy, which can be estimated by LZ and other methods such as BDM (discussed below), provide a more direct link with the subjective structure of experience [3, 65].

After applying the algorithm to a string of length *n*, we obtain a set of words in a dictionary of length *c*(*n*) that can be thought of as a first complexity metric [66]. The description length of the sequence encoded by LZ can be approximated by the number of words seen times the number of bits needed to identify a word, and, in general, ℓ ≈ *c*(*n*) log_2_ *c*(*n*) ≲ *n*. We note that LZ provides an upper bound on algorithmic complexity, but, given its reduced programming repertoire (LZ is the Kolmogorov complexity computed with a limited set of programs that only allow copy and insertion in strings [67]), it fails, in general, to effectively compress random-looking data generated by simple, but highly recursive programs, e.g., an image of the Mandelbrot set (deep programs [3, 68]). As a simple example of these limitations, consider a string concatenated with its bit-flipped, “time-reversed”, or dilated version. Such simple algorithmic manipulations will not be detected and exploited by LZ. Despite these limitations, LZ can be useful for studying the complexity of data in an entropic sense. To use LZW, we first transform numeric data into a list of symbols. Here we binarize the data to reduce the length of strings needed for the algorithm to stabilize. A reasonable strategy in the study of algorithmic information content in data is to preserve as much information as possible in the resulting transformed string. In this sense, using methods that maximize the entropy of the resulting series is recommended. Here, as discussed above, we use the median of the data as the threshold.

Two associated metrics derived from LZ compression are commonly used: *c*(*n*) and *l*. Of the two, the latter is more closely related to Kolmogorov complexity or description length. A natural way to normalize this metric is to divide the description length by the original string length *n*, 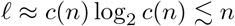, with units of bits per character. Here, we analyze synthetic lattice data generated by the Ising model by computing *ρ*_0_ for each lattice realization. We also analyze *ρ*_0_ by concatenating them for a global measure (with real data). The former allows for a plot of the mean and standard deviation of the measure with synthetic data generated using the Metropolis algorithm as a function of temperature (Figure 3).

To calculate a global measure of LZ complexity using the experimental data, we first concatenated the data from all subjects, conditions, and sessions then flattened it along the spatial dimension first and then time. Secondly, we compressed the flat binarized data with the LZ algorithm and saved the output “archetype dictionary”, which was used as the initial dictionary to compress the data from all the subjects in the two conditions, LSD and placebo (comparison by condition), and also per subject (comparison by condition and subject). The archetype dictionary was built by concatenating first the LSD data for all subjects and sessions and then the placebo data because we were forced to make a choice when flattening the array. However, we checked whether concatenating the placebo data before the LSD changed the results, and the difference was negligible. For details and code used to compute LZ complexity—we use the Lempel-Ziv-Welch (LZW) variant—see [55, 69].

#### Block decomposition method

The Block Decomposition Method (BDM) [70] combines the action of two universally used complexity measures at different scales by dividing data into smaller pieces for which the halting problem involved can be partially circumvented, in exchange for a huge calculation based upon the concept of algorithmic probability and extending the power of the so-called Coding Theorem Method (CTM). The calculation, however, has been pre-computed and hence can be reused in future applications by exchanging time for memory in the population of a pre-computed look-up table for small pieces. BDM is used to account for both statistical regularities and algorithmic ones and is sensitive enough to capture small changes in the complexity while invariant to different descriptions of the same object. It, therefore, complements the use of lossless compression algorithms to calculate an upper bound of algorithmic complexity. We carry out the same analysis as for *ρ*_0_ described above, i.e., both on synthetic and experimental data.

#### Ising spin model

Abstract frameworks from statistical physics can shed light on understanding emerging phenomena in large networks, such as phase transitions in systems where nodes—neurons or cortical columns here—interchange information under the assumptions of the maximum entropy principle [49, 50, 71]. The description of systems with many degrees of freedom can be summarized by coarse-graining variables (describing macrostates), which introduces statistics into modeling. At the so-called critical points, observable quantities such as diverging correlation length or susceptibility to external perturbations reveal singularities in the limit as the number of degrees of freedom goes to infinity. At these transitions, from order to disorder, there is a loss of sense of scale, with fractal properties in energy and information flow.

In this context, elements such as neurons, columns, or brain regions are modeled by spins (i.e., with two states, up or down, on or off) with pair interactions. The emerging statistical properties of large networks of these elements are studied under different conditions (temperature or an external magnetic or electric field). The prototypical simplest system in this context is the classical 2D Ising model, which features nearest-neighbor interactions and a phase transition. This model has been shown to be universal, i.e., that all the physics of every classical spin model (with more general types of interactions) can be reproduced in the low-energy sector of certain “universal” models such as the 2D Ising model [72]. This fact reflects the intrinsic computational power of near-neighbor interactions. In fact, Ising models have been shown to be universally complete [73, 74], with a map between any given logic circuit to the ground states of some 2D Ising model Hamiltonian. On the other hand, the fact the brain exhibits characteristics of criticality that may be modeled by systems such as the Ising model is now well established, with ideas that go back to pioneers such as Turing [75], Bak [4, 76], and Hopfield [77]. There is further evidence that the dynamics of the healthy brain occupy a sub-critical zone ([78], and see also [32] and references therein).

The Ising spin model used here arises naturally from model fitting of the binarized data (discussed in the next section) and is slightly more general than the original one. It is essentially the Sherringon-Kirkpatrick spinglass model [79] with an external field and allows for arbitrary pair, arbitrary sign interactions. It is defined by the energy or Hamiltonian of the lattice of *N* spins,

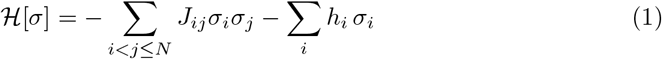

where *σ_i_* denotes the orientation of each spin in the lattice (±1), *J_ij_* is the coupling matrix or Ising connectivity (with pairs counted once), and *h_i_* is an external magnetic field applied independently at each site. In the context of analysis of parcellated BOLD data, *σ_i_* in this formalism represents the binarized state of each brain parcel and is equal to +1 (— 1) when the parcel is active (inactive), *h_i_* is a parameter that modulates the mean activity in a single parcel, and 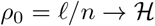 is a parameter that accounts for the interactions between parcels *i* and *j* (correlations), and which can be positive or negative.

#### Maximum entropy principle derivation of Ising model

The (spinglass) Ising model, which was originally proposed to describe the physics of ferromagnetic materials, can be seen to arise naturally from the Maximum Entropy Principle (MEP) [49]. The MEP is used to find an appropriate probability distribution given some constraints, e.g., derived from data. It states that the probability of finding a given spin configuration ***σ*** can be inferred from a simplicity criterion: given some data-derived constraints—which here will be that the mean spin and spin correlation values of the model should match that of the data—the probability function we should use is the one that fits the constraints but has otherwise maximal Shannon entropy. It turns out that the probability distribution function of observing a given spin configuration ***σ*** (bold symbols indicate an array) in thermal equilibrium (at unit temperature without loss of generality) is given by the Boltzmann distribution [8]

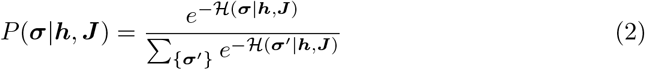

with 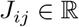 given in Equation 1, and where 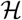 indicates a sum over all the 2^N^ possible spin configurations.

#### Estimation of archetype parameters using maximum pseudo-likelihood

To create the global model archetype, we concatenate the binarized data from the fifteen participants and the two conditions (a total of four sessions) to generate a single dataset. We similarly produce condition (LSD and placebo) archetypes by concatenating the data in each condition.

We estimate the model parameters **J** and **h** using an approximation to maximum likelihood as described in *Ezaki et al*. [8]. Briefly, we find the parameters that maximize the probability of observing the data given the model,

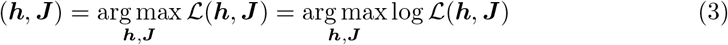

with

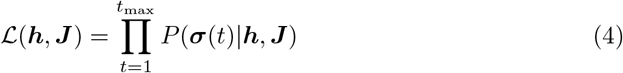

The likelihood maximum can be found by gradient ascent, with

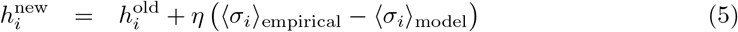

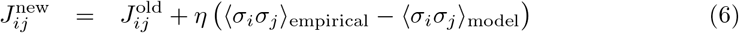

where the old/new superscripts refer to the values before and after updating, and with *η* a small positive constant.

When the number of nodes in the system is large, the calculation of the likelihood from the model because computationally intractable. For this reason, it is approximated by the pseudo-likelihood, a mean-field approximation that approaches the true likelihood as the number of time points goes to infinity [8, 80],

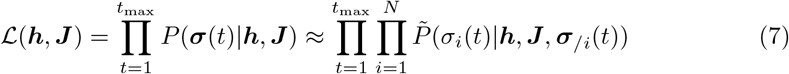

with 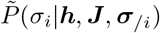 the modeled probability distribution for a given spin given the states of all the others, a quantity much easier to compute,

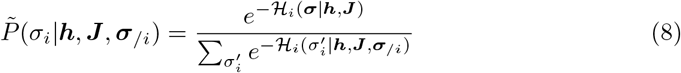

with

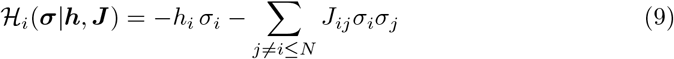

Using this approximation, the gradient ascent rule becomes

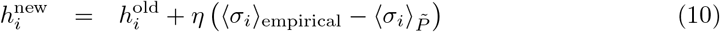

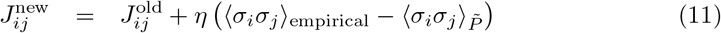

where 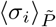 and 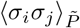 are the 1-point and 2-point correlation functions with respect to distribution 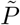 [8],

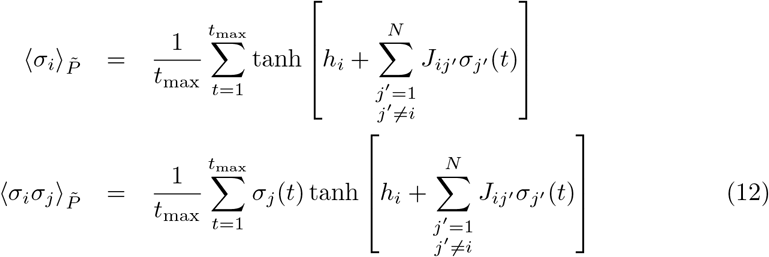

#### Personalization of model—individualized temperature

With ***h*** and ***J*** fixed in the archetype using the entire dataset, we adapt the archetype model for each subject and condition by changing the model temperature *T* = 1/*β*, that is, by writing

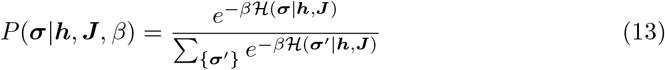

In this case, the gradient ascent algorithm becomes

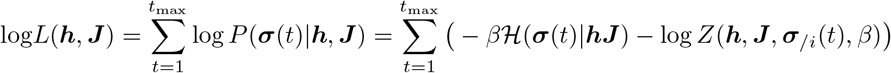

with a fixed point at 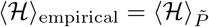. This, as in the prior equations, can be seen by taking the derivative of the approximate log-likelihood

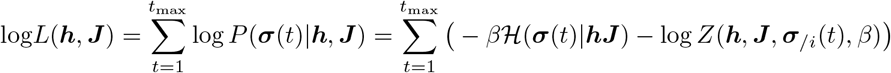

with respect to the inverse temperature *β*, with the partition function, *Z*, given by

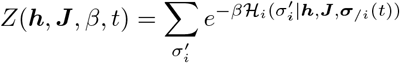

The first term becomes the empirical average of the Hamiltonian, and the second the (data dependent) model mean (both up to a constant since we are not dividing by the number of measurements). It can be computed from Equation 12 and from

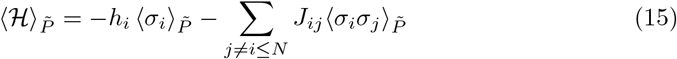

#### Metropolis algorithm and Ising model observables

The Metropolis algorithm is used to compute the observables of the Ising model built from the experimental data. A loop is performed where a random spin is chosen and flipped with a probability 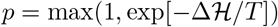, where 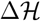 is the change in energy the flipped spin would cause in comparison to the previous configuration and *T* is the system temperature. This process provides the means to sample the system probability distribution at a given temperature. The method is applied for a sufficient number of iterations to reach a steady state, and the macroscopic variables are obtained as a mean across iterations over a large number of configurations in steady state. To evaluate 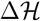, we rewrite the energy of the system (Hamiltonian) as

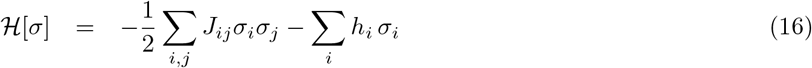

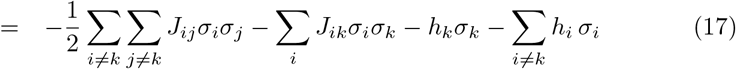

where in the second line, we separate out the contribution to the energy from spin *σ_k_* (which receives two terms from *J*, which is symmetric). The energy associated with the *k*-th spin is

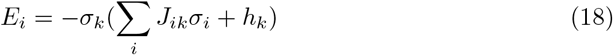

After a flip of this spin, its energy contribution becomes

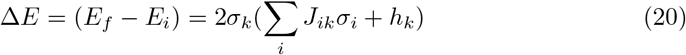

and

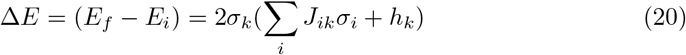

The main global observable is the lattice average magnetization, *M*, over the spin lattice, *S*, of *N* spin configurations, *σ_i_*,

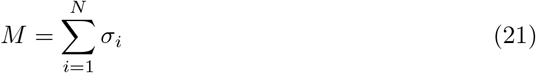

Then, the magnetic susceptibility, *χ*, and heat capacity, *C_v_*, can be computed as follows. Let us apply a uniform external field *h*,

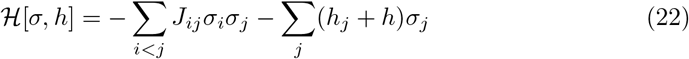

and let

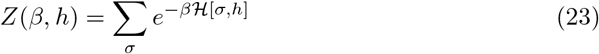

be the partition function. The average magnetization, 〈*M*〉, is the ensemble average of lattice magnetization,

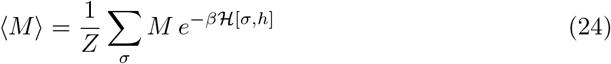

Then it can be verified that (see, e.g., [81])

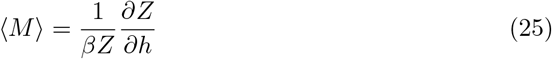

Following similar considerations (with *T* = 1/*β*), the global susceptibility, *χ*, and the system heat capacity, *C_v_*, are given by

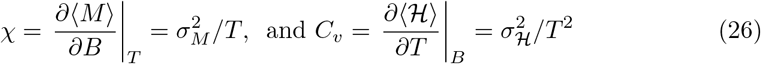

where the standard deviation computation is taken over the ensemble. We can extend the definition of global susceptibility (obtained from average quantities of the system) to local susceptibility, *χ_n_*, (specific to each spin or brain parcel). Let

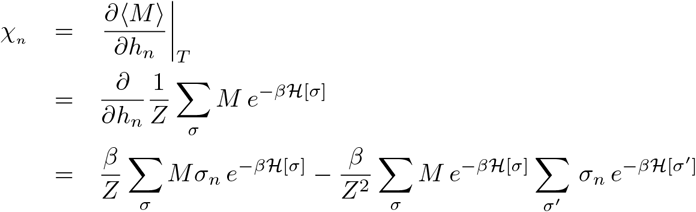

so

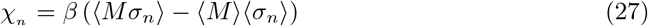

The global and local node susceptibility quantify, respectively, the sensitivity of the lattice state (magnetization) to an external uniform or local perturbation (represented by a magnetic field). They are interesting quantities in the context of brain dynamics from various perspectives, including the effects of external perturbations such as sensory inputs or brain stimulation. Similarly, the link susceptibility

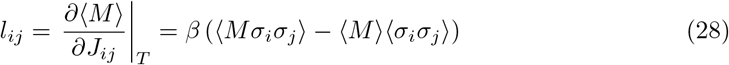

provides information about the sensitivity to additive changes to a particular link in the system. To understand the impact of changes of scale of a particular link (*J_ij_* → *J_ij_* + *αJ_ij_*), we define the corresponding scaled quantity Lj = *ljJj*.

To smooth the results and deal with slow lattice fluctuations at cold temperatures in finite-size systems (the system is bistable for the finite case), we work with modified variables to compute the magnetization and susceptibility statistics using Metropolis-generated data. Selection of one of the metastable branches is achieved by flipping, as needed at each sample point, the entire lattice so that the magnetization is positive, *σ_n_* → *σ_n_* sign(*M*). E.g., *M* → |*M*| = *M* sign(*M*) for the evaluation of magnetization and global susceptibility (v. Figure 2). Even *n*-point correlation functions such as 〈*Mσ_n_*〉 are invariant. The results away from the critical temperature differ by a constant from the ideal one (the effect is one of smoothing near the critical point) [82].

### Statistical analysis

For the analysis of the Ising models, we performed a paired Wilcoxon test of the temperature estimates. To further test whether the observed increase in temperature under LSD relative to placebo was specific to the architecture parameters *h* and *J*, we conducted a non-parametric permutation test on the mean temperature difference [83]. In each of one thousand permutations, we randomly shuffled the parcellation labels and hence the *h* and *J* components of the archetype parameters (see Equation 3). These new set of permuted archetypes (*h_p_, J_p_*) were then used to fit the system temperature of each participant and condition (see Equation 13). The null hypothesis is that the measured BOLD activity patterns are drawn from the same probability distribution, regardless of the drug manipulation in which it was recorded. Once the personalized *β_p_* estimates were obtained, we performed the temperature contrast between LSD and placebo. This was carried out by computing a one-sample t-test for every permutation. The level of statistical significance was estimated by the fraction of t-values derived for every random shuffle that was higher than the observed t-values obtained from our original temperature contrast obtained from the non-permuted h and J. Similarly, we performed a paired one-tailed Wilcoxon test with the LZW and BDM complexity estimates.

To compare archetypes, we used paired t-tests (Wilcoxon signed-rank tests provided very similar results). We used Pearson correlation to study correlations of connectivity matrices, metrics, and receptor maps.

### dMRI tractography

We will find it useful to compare Ising connectivity with structural connectivity obtained from dMRI tractography in a separate cohort as described in Cabral et al., 2014 [84]. Briefly, the data was obtained from 21 healthy, normal participants (11 males and 10 females, age: 22-45 years) different from the healthy subjects in the LSD cohort. All scans were performed on the same Philips Achieve 1.5 Tesla Magnet. Diffusion MRI was acquired by using a single-shot echo-planar imaging-based sequence with coverage of the whole brain with 33 optimal nonlinear diffusion gradient directions (b = 1200 s/mm2) and 1 non-diffusion weighted volume (b = 0), repetition time (TR) = 9390 ms; echo time (TE) = 65 ms. T1-weighted structural images with a three-dimensional ‘FLASH’ sequence (TR = 12 ms, TE = 5.6 ms, flip angle = 19°, with elliptical sampling of k-space, giving a voxel size of 1×1×1 mm in 5.05 min) were also acquired. The AAL template was used to parcellate the brain into 90 regions (45 for each hemisphere), which define the network nodes. For each participant, parcellation was conducted in the diffusion-MRI native space. The b0 image in diffusion-MRI space was linearly co-registered to the T1-weighted structural image using the Flirt tool (FMRIB, Oxford). The transformed T1-weighted image was then mapped to the T1 template of ICBM152 in MNI space, inversed and further applied to warp the AAL mask from MNI space to the diffusion-MRI native space. Interpolation using the nearest-neighbor method ensured the preservation of labeling values. The links between nodes were weighted in proportion to the number of white matter tracts detected between brain areas. The processing of the diffusion-MRI data was performed using the Fdt toolbox in FSL (www.fmrib.ox.ac.uk/fsl, FMRIB). Pre-processing involved the co-registration of the diffusion-weighted images to a reference volume using an affine transformation to correct head motion and eddy current-induced image distortion. Subsequently, the local probability distribution of fiber direction at each voxel was estimated. The *probtrackx* algorithm was used, allowing for automatic estimation of two fiber directions within each voxel, which can significantly improve the tracking sensitivity of non-dominant fiber populations in the human brain. For each voxel in the brain, probabilistic tractography was used to sample 5000 streamline fibers passing through that voxel. The connectivity probability from voxel *i* to another voxel *j* was defined by the proportion of fibers passing through voxel *i* that reach voxel *j*. This was then extended from the voxel level to the region (parcel) level. The connectivity *C_np_* from region n to region *p* is calculated as the number of fibers passing through any voxel in parcel *n* that connect to any voxel in parcel *p*, divided by 5000×*N*, where *N* is the number of voxels in parcel *n*. For each brain region, the connectivity to each of the other 89 regions was calculated. Since the connectivity from *n* to *p* is not necessarily equal to that from *p* to *n* but highly correlated for all subjects, the undirected connectivity *C_np_* between regions *n* and *p* was defined by averaging the two.

## Results

### Ising connectome vs. dMRI connectome

The first step in our analysis is to identify the parameters *h* and *J* that could best fit the fMRI data from all subjects in both conditions—which leads to what we call the ‘archetype model.’ Additionally, in order to check for network changes due to the effect of LSD, we also estimated the parameters *h* and *J* that could best reproduce the data from the LSD and placebo conditions separately. Gradient descent resulted in very good fits of the functional connectivity data in each condition (*r* ~ 1, see Figure SI.11).

Figure 1 (top) provides the resulting connectivity matrix (i.e., the values of *J_ij_*) estimated from all the data and also the changes in connectivity between the LSD and placebo conditions. We can observe that structural intrahemispheric connectivity in the Ising model is mostly positive. In contrast, interhemispheric connectivity is mostly negative and weaker except for homotopic links (strong and positive).

**Fig 1.**
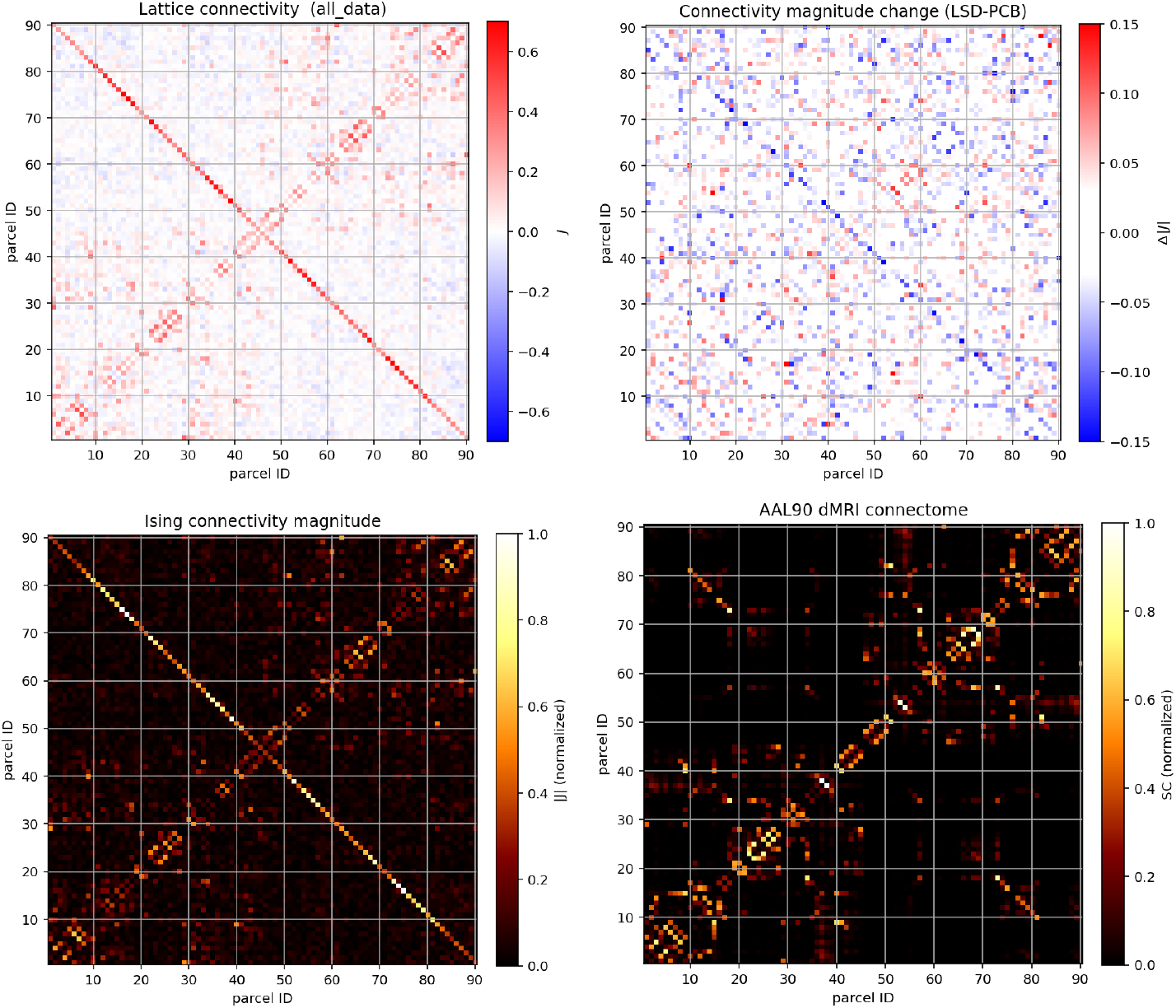
Archetype connectivity. **Top left:** global archetype (combining all data) Ising connectivity. The down-trending diagonal values reflect strong inter-hemispheric homotopic connectivity. **Top right:** difference in connectivity magnitude between the placebo and LSD archetypes, with a visible loss in homotopic connectivity (see also figures in SI Section F). **Bottom left:** unsigned global archetype Ising connectivity. **Bottom right:** AAL connectome (from [85]).

Figure 1 (bottom) provides an unsigned version of the global archetype connectivity matrix *J* and, for reference, the structural connectome (obtained with dMRI tractography from a separate cohort as described in Cabral et al., 2014 [84]—see Methods). We can see that the two are very similar (*r* = 0.60). The correlation between the two intrahemispheric submatrices is higher (*r* = 0.70), while for the interhemispheric ones is strong (*r* = 0.54) but mostly accounted for by the homotopic diagonal (removing it reduces *r* to 0.14).

### Differences between LSD and PCB archetype connectivities

To analyze the archetypes, we introduce further metrics. We define the unsigned Global Connectivity (GC) as the array resulting from summing the rows of the magnitude (absolute value) of the connectivity adjacency matrix,

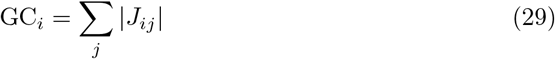

We also introduce the interhemispheric and intrahemispheric submatrices, *J*^inter^ and *J*^intra^, where only the respective links are included (others set to zero), and their respective global connectivities GC^inter^ and GC^intra^. Finally, we also use the notion of homotopic connectivity (HC). In our parcellation, we label homologous areas symmetrically (see Appendix A), and

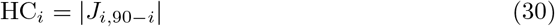

The change in link strength between |*J*_LSD_| and |*J*_placebo_| was significant (statistic=-4.4, p=1.09×10^-5^, treating the array elements in the two archetypes as samples). No significant changes were found in the signed version of global connectivity between the two archetypes, but they were strong in the unsigned version of this quantity (GC) (t-test statistic=-7.6, p=3.1 × 10^-11^, Wilcoxon p=2.3 × 10^-9^). The changes in *h* exhibited a trend to weaker external uniform drive (t-test statistic=-1.54, p=0.13) on the system. The main effect, which can be observed in Figure 1, was a robust decrease in homotopic connectivity across parcels (connectivity between homologous areas in the two hemispheres, HC) in the LSD condition (t-test statistic=—7.8, p=7.6 × 10^-10^, Wilcoxon p=1.1 × 10^-7^). These changes at the level of the two archetypes reflect differences in the total homotopic FC for each subject across the two conditions (paired Wilcoxon statistic=-4.38, p=6×10^-4^). In more detail, interhemispheric (t-test statistic=—6.9, p=6.1 × 10^-10^, Wilcoxon p=8.2 × 10^-9^) but not so much intrahemispheric GC (t-test statistic=—2.9, p=0.005, Wilcoxon p=0.003) was significantly reduced in the LSD condition. The decrease in homotopic connectivity scaled with homotopic connectivity estimated in the placebo condition (*r* = —0.4, p=1.1 × 10^-5^).

### Archetype connectomes and 5-HT_2A_ distribution

As the LSD effects should be related to the density of the 5-HT_2A_ receptors [1, 86], we analyzed the correlation between the receptor density distribution, Ising connectivity, and the changes in the archetype parameters obtained separately from the LSD and placebo data. The 5-HT_2A_ receptor map in AAL parcellation was obtained as in Deco et al. (2018) [86]. We found a weak negative correlation between homotopic connectivity and receptor density (*r* =-0.27, p= 0.01) and a positive correlation between GC^intra^ and receptor density (*r*=0.37, p=3×10^-4^), both especially evident in the LSD condition. For the overall GC and receptor density, the correlation was weaker (*r*=0.24, p=0.02 with data from both conditions). No correlation was observed between receptor density and GC^inter^. Finally, we did not observe correlations between receptor density and changes in GC or homotopic connectivity.

### Archetype phase diagrams

To find the critical temperature of the archetype model, we explored its phase space by calculating key quantities belonging to the Ising formalism and related measures of signal and algorithmic complexity. Figures 2 and 3 provide, respectively, the local susceptibility and local magnetization at each node in the lattice, and the global magnetization, susceptibility, energy, heat capacity, LZW, and BDM complexity estimates and their standard deviation, as a function of temperature. As can be observed, the critical point (i.e., the peak of the standard susceptibility) is well below *T* =1, the nominal temperature of the archetype model (which corresponds to the global fit) and of the individualized models. This indicates that archetype and individualized models are all in the paramagnetic, supercritical phase. Susceptibility, heat capacity, and complexity peak at approximately the same temperature (*T* ~ 0.7-0.8). Finally, Figure 2 displays the peak susceptibility of nodes in the parcellation and the temperatures at which they occur.

**Fig 2.**
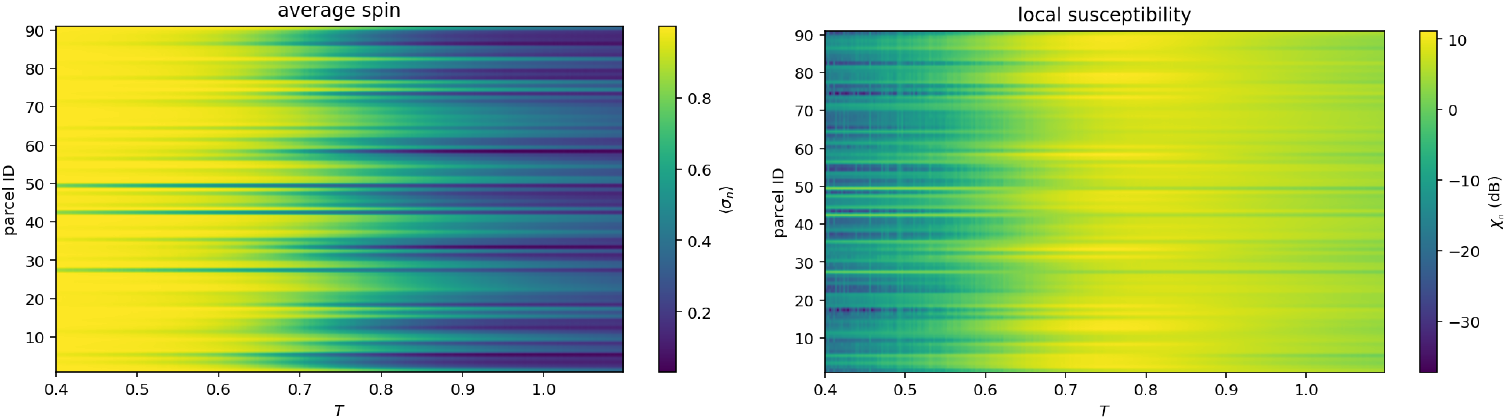
Local magnetization and susceptibility from Metropolis simulations with the global archetype model. Local magnetization and susceptibility (i.e., specific to each spin or brain parcel) of the archetype model as a function of temperature.

**Fig 3.**
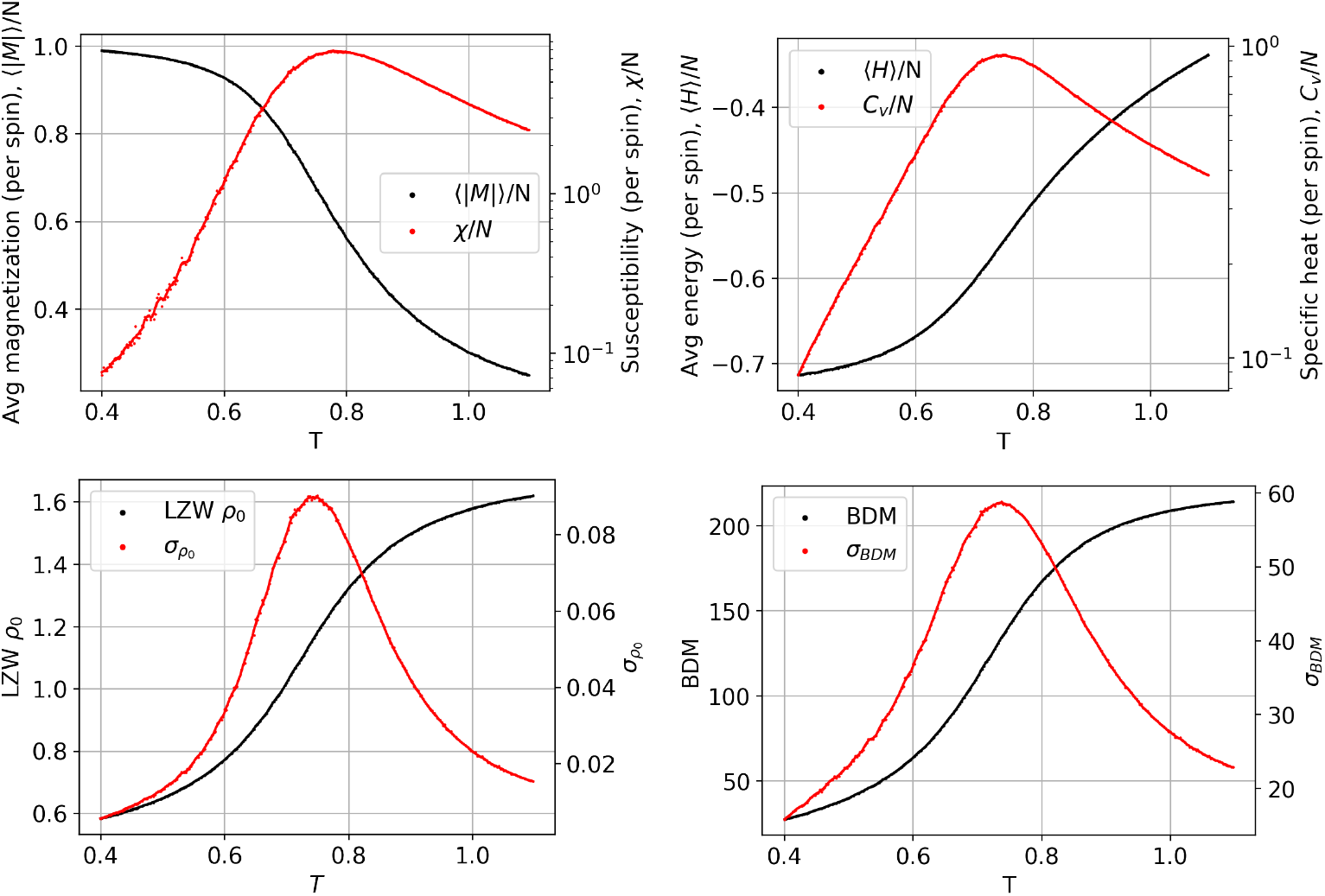
Phase diagram for global archetype model. Top: Global susceptibility and magnetization (left), energy and heat capacity (right). Bottom: LZW complexity (*ρ*_0_) and its standard deviation (left), and BDM complexity and its standard deviation (right), for the archetype model as a function of temperature (the Metropolis algorithm has been run for 10^8^ spin flip tests for each temperature to obtain these plots).

### Individual temperature shift with LSD

The next step in our analysis was to analyze the personalized temperatures of each subject during the LSD and placebo conditions. These temperatures can be compared to the system’s critical temperature to locate where the temperatures of the two conditions and different subjects sit in the archetype phase space. Figure 4 displays, on the top, personalized temperatures of the fifteen subjects in the LSD and placebo conditions and the temperature difference between conditions for each subject.

**Fig 4.**
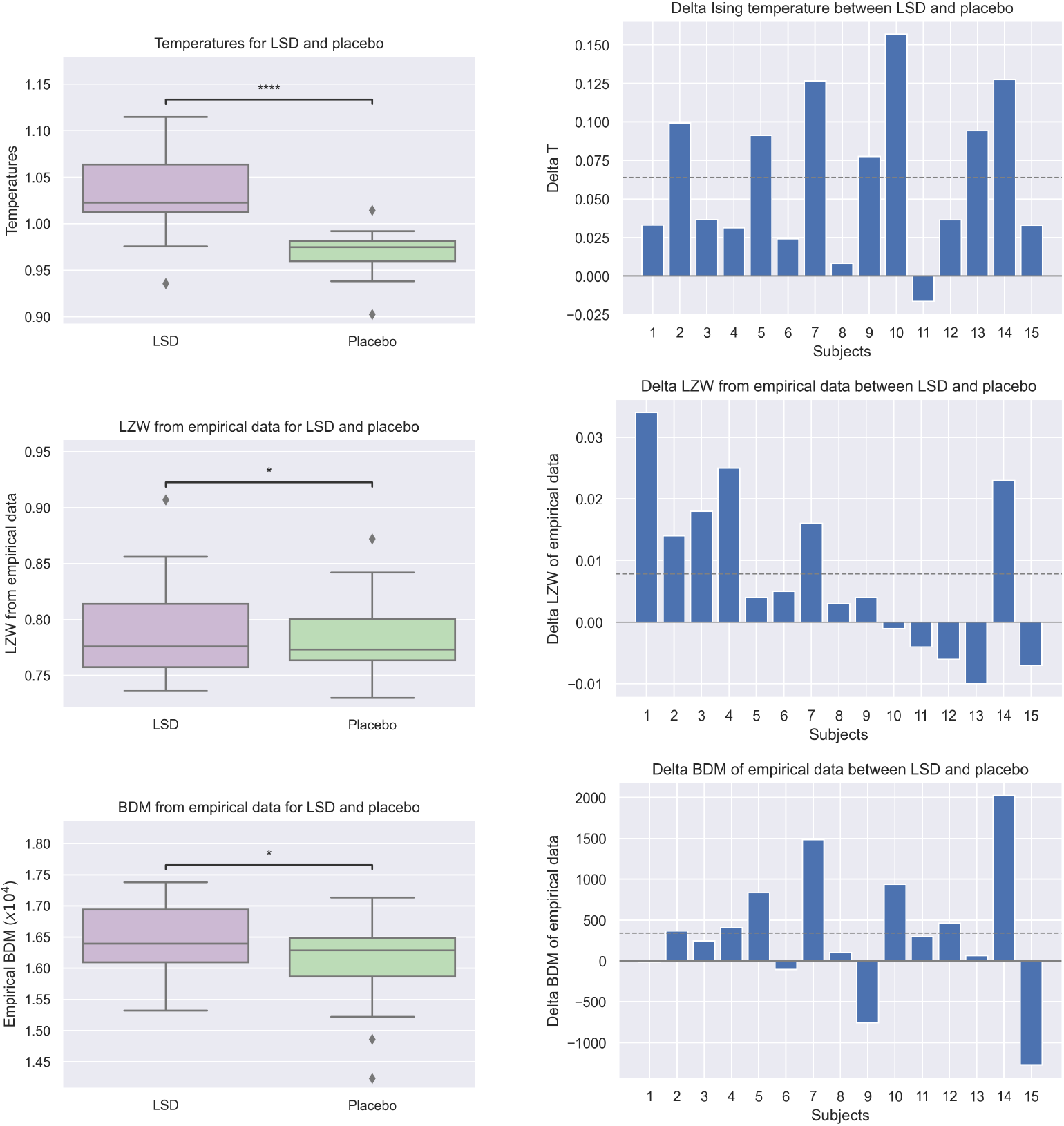
Individual shift of Ising temperature (top), LZW (middle), and BDM (bottom) complexity estimates with LSD using empirical data. Left: Box plots of the metrics of 15 subjects under the effect of LSD and placebo. Right: Bar plot of the metrics differences between LSD and placebo for each subject. The dashed line represents the mean metric. (One star: *p* < 0.05, four stars: *p* < 0.0001)

Temperatures in the LSD state were found to be substantially higher than in placebo (6.7% ± 5.1%; mean ± standard deviation), with this increase being statistically significant (two-tailed t-test *p* = 6.1 × 10^-5^; Cohen’s *d* = 1.35, one-tailed Wilcoxon test *p* = 9 × 10^-5^). The non-parametric permutation test shuffling ROI labels in the model to test sensitivity to archetype parameters confirmed these were significant results (*p* =1 × 10^-3^).

### LZW and BDM complexity from empirical and model data

We also extracted features related to the algorithmic complexity both from the empirical fMRI data and the synthetic data generated with the model with the Metropolis algorithm. We would expect these features to show a similar trend to the one obtained with the Ising temperatures. Figures 4 and 5 display the individual LZW and BDM complexity of 15 subjects under the effect of LSD and placebo (left) and the difference in LZW and BDM complexity between the LSD and placebo conditions for each subject (right), calculated respectively using experimental and simulated data generated by the personalized Ising models.

**Fig 5.**
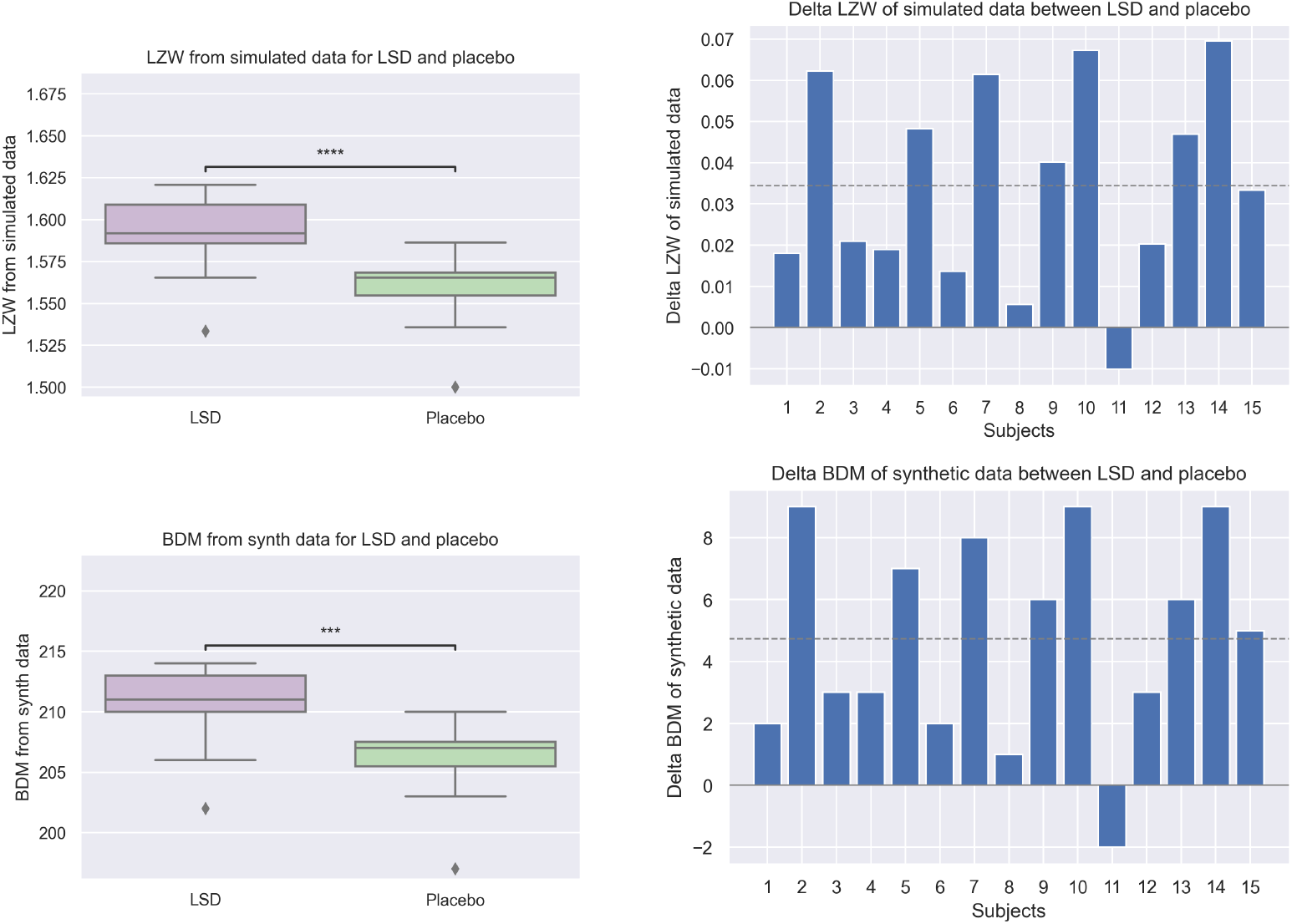
Individual shift of LZW (top) and BDM (bottom) complexity estimates with LSD using synthetic data. Left: Box plots of the metrics of 15 subjects under the effect of LSD and placebo. Right: Bar plot of the metrics differences between LSD and placebo for each subject. The dashed line represents the mean metric. (Three stars: *p* < 0.001, four stars: *p* < 0.0001)

The LZW complexity in the LSD state from the data was found to be higher than in the placebo condition for 2/3 of the subjects (Figure 4, middle). The mean and median relative LZW complexity shift with LSD were respectively 0.008 and 0.005, with a standard deviation of 0.013. The paired one-tailed Wilcoxon test with the LZW complexity estimates resulted in *p* = 0.04 for the Wilcoxon test (*p* = 0.04 for the t-test). The probability of finding this or a larger increase of *ρ*_0_ with LSD ingestion was estimated at *p* < 0. 04 (one-tailed Wilcoxon test).

As shown in Figure 4 (bottom), the BDM complexity in the LSD state inferred directly from the data was higher for more than 70% of subjects. The mean and median BDM complexity shift with LSD were respectively 338 and 298, with a standard deviation of 804. Also, the BDM complexity from empirical data correlated with the condition (*p* = 0.04, one-tailed Wilcoxon test).

As can be observed in Figure 5, the increase in LZW and BDM complexity in the LSD state, using synthetic data generated by the personalized models, was statistically significant (one-tailed Wilcoxon test *p* = 9 × 10^-5^ and *p* = 2 × 10^-4^, respectively).

We also analyzed the correlation between the features extracted from the data and from the model. A summary of the statistics of the different metrics and the Pearson correlation coefficients between them is given in Tables 1 and 2, whereas the correlation scatter plots can be found in Figure SI.16 in Appendix J. The difference in LZW and BDM complexity from the model correlates strongly with the delta Ising temperature (*r*(13) = 0.97, *p* = 3.46 × 10^-9^ for LZW and *r*(13) = 0.95, *p* =1.1 × 10^-7^ for BDM). The changes in LZW from experimental data do not display a significant relationship with the changes in Ising temperature (*r*(13) = 0.06, *p* = 0. 83), nor with the delta LZW complexity from synthetic data (*r*(13) = 0.10, *p* = 0.73), presumably due to the short length of the time series and the limited number of subjects. The difference in BDM complexity from the experimental data correlates with the delta Ising temperature (*r*(13) = 0.56, *p* = 0.03), whereas it does not show a strong statistical relationship with the delta BDM from the model (*r*(13) = 0.42, *p* = 0.12), with the delta LZW from the data (*r*(13) = 0.35, *p* = 0.19), and with the delta LZW complexity from the model (*r*(13) =0.48, *p* = 0.07).

**Table 1.** Summary statistics of different metrics. All metrics refer to the change of variable (LSD-Placebo).

|  | Median (%) | Mean (%) | t-score | p-value (one-tailed) |
| --- | --- | --- | --- | --- |
| Ising $T$ | 3.76 | 6.61 | 4.93 | $9 \times 10^{-5}$ |
| LZW data | 0.50 | 1.00 | 2.30 | 0.04 |
| LZW synth | 2.13 | 2.21 | 5.46 | $9 \times 10^{-5}$ |
| BDM data | 1.83 | 2.10 | 1.63 | 0.04 |
| BDM synth | 2.51 | 2.28 | 5.18 | $2 \times 10^{-4}$ |

**Table 2.** Pearson correlation between different metrics.

| Pearson cc | Ising $T$ | LZW data | LZW synth | BDM data | BDM synth |
| --- | --- | --- | --- | --- | --- |
| Ising $T$ | | 0.06 | 0.97**** | 0.56* | 0.95**** |
| LZW data |  |  | 0.10 | 0.35 | 0.08 |
| LZW synth |  |  |  | 0.48 | 0.99**** |
| BDM data |  |  |  |  | 0.42 |

### Correlation between questionnaire scores and data features

In the LSD trial, participants were asked to answer two types of questionnaires to give feedback on their psychedelic experience. They completed VAS-style ratings via button-press and a digital display screen after each scan and the 11-factor altered states of consciousness (ASC) questionnaire at the end of each dosing day [33]. The most important ASC scores reported by the authors were elementary imagery (“I saw regular patterns in complete darkness or with closed eyes.”, “I saw colors before me in total darkness or with closed eyes.”, “I saw lights or flashes of light in total darkness or with closed eyes.”), complex imagery (“I saw scenes rolling by in total darkness or with my eyes closed.”, “I could see pictures from my past or fantasy extremely clearly.”, “My imagination was extremely vivid.”), and audio-video synesthesia (“Noises seemed to influence what I saw.”, “The shapes of things seemed to change by sounds and noises.”, and “The colors of things seemed to be changed by sounds and noises.”).

We calculated the questionnaire scores difference between the LSD and placebo sessions for each category. Then, we computed the average scores of each questionnaire to have a single measure of the overall intensity of the subjective experience. We checked the Pearson correlation between the subjects’ questionnaire scores and the features extracted from the data, i.e., Ising temperature, LZW, and BDM from experimental data.

Tables SI.16 and SI.16 in Appendix K provide the questionnaire scores for all subjects, their average, and the Pearson correlation coefficients between the scores and the features extracted from the data.

No strong statistical relationships were seen between the subjective reports and the features. However, trends were observed between complex imagery and Ising temperatures (*r* = 0.42) and elementary imagery and LZW complexity (*r* = 0.48), both obtained from the ASC questionnaire. From the VAS-style ratings, trends were observed between emotional arousal and all three features (positive trends for Ising temperatures and BDM complexity, whereas negative ones with LZW complexity).

## Discussion

We have developed here a framework for the generation and analysis of personalized Ising spin (spinglass) models starting from group archetypes. We discuss our results next.

### Ising and structural connectome

A first finding from fitting the (maximum entropy) Ising model to functional BOLD data is that it is possible to recover, to a good extent, the underlying structural connectome as inferred from dMRI data. In fact, what is recovered is a signed approximation to the structural connectome, which may be useful for whole-brain modeling using more sophisticated models such as neural mass models where it is not easy to decide where long-range connectivity should be inhibitory. In addition to negative couplings, the biggest discrepancy between the two connectomes is the homotopic component of connectivity (v. Figure SI.8), both of which reflect features of the FC (v. Figure SI.12 and the figures in Section SI G). This is not surprising since one of the most interesting differences between functional and structural connectomes is precisely the homotopic component, which is especially salient in resting-state fMRI functional connectivity [87] despite not being structurally as strong as detected by dMRI tractography [88, 89]. This discrepancy suggests that homotopic functional connectivity may arise from indirect links through other areas, e.g., by a synchronizing role of subcortical areas such as the thalamic nuclei, achieved by interacting symmetrically with both hemispheres [90], or from direct neuromodulation of structural elements (the tractogram provides an estimation of the number of fibers connecting parcels, but it does not provide direct information about the number of synapses associated with the fibers or the connectivity state as induced by neuromodulatory agents, for example). In either case, the Ising model must reproduce homotopic functional connectivity clearly present in the resting state data (parcel state correlations), and its structural elements (*J*) reflect this. An additional observation is that the Ising connectome was mostly dominated by positive intrahemispheric and homotopic links. Interhemispheric connectivity was otherwise weak and mostly negative in our dataset.

### Ising archetypes for LSD and placebo conditions. 5-HT_2A_ distribution

With regard to the archetypes built separately in each condition, the main difference between the global LSD and placebo archetypes was a decrease in connectivity under LSD, mostly associated with homotopic links. We observed the largest losses in homotopic connectivity in the calcarine, hippocampus, thalamus, and fusiform parcels (see Figures SI.2 and SI.6). The increase of disorder under the LSD condition, as quantified by Ising temperature or complexity metrics, was thus associated with a loosening of interhemispheric connectivity driven by a loss of homotopic connections (Figure 1, right). The decrease in homotopic connectivity was, in turn, proportional to the placebo condition archetype homotopic connectivity. Our results suggest a transient reduction, induced by LSD, of Ising structural homotopic connectivity, where strong homotopic connections see the largest decrease in strength. Such a reduction has a strong impact on the system because the Ising connectome can be seen to consist of two networks (one per hemisphere) with weaker interhemispheric connections that are dominated by homotopic links providing “long-range” connectivity. The effect of reducing these connections can be simulated in the Ising formalism (v. Figure SI.14 in Appendix I). E.g., reducing homotopic connectivity in the placebo archetype by twenty percent leads to a decrease of its critical temperature of ten percent (from *T* = 0.82 to 0.74), making the system at nominal temperature (*T* = 1) more disordered (reducing an equal number of random connections has a smaller impact with a mean *T* = 0.80 with a standard deviation of 0.02). Another way to study the sensitivity of the system to changes in its connections is through the concept of linked susceptibility as defined above (*L_ij_*). In particular, this displays increased values along the homotopic line (Figure SI.15). With regard to the potential mechanism for these changes, we found that the density of 5-HT_2*A*_ receptors was positively correlated with the strength of intrahemispheric global connectivity and negatively correlated with the homotopic connectivity of each parcel, suggesting that the relative functional weight of homotopic links may be reduced by the increased intrahemispheric 5-HT_2*A*_ agonistic effects of LSD.

### Global archetype critical point and local susceptibility

We found it especially useful to generate a global archetype (v. Figure SI.7) assimilating the data from all participants and conditions (see also Figure SI.5 for the empirical FC of all the data). This is convenient first because of the relative scarcity of data produced by BOLD fMRI sequences and, second, because it provides a common framework for comparison of individualized models (in this case, characterized by their temperatures relative to the archetype). The archetype generates behavior as a function of temperature similar to the standard, nearest-neighbor Ising model, with a clearly defined critical point. However, unlike the original Ising model, this model does not display translational symmetry because the model parameters (*J_ij_* and *h_i_*) vary with each parcel. As a consequence, some of the nodes of the archetype model display higher sensitivity to perturbations than others (see figures in SI Section C). The brain parcels corresponding to these nodes are natural areas for further study of the effects of stimulation (invasive or non-invasive) in more sophisticated computational brain models [91, 92].

### Personalized temperatures and complexity

With regard to the personalized temperatures, we found a strong correlation between individualized temperature and condition. The individualized temperatures derived from BOLD fMRI data almost uniformly increased with the LSD condition relative to the placebo. Statistically speaking, the results are very robust. Moreover, both the archetype temperature (*T* = 1) and the individualized ones for both conditions were found to be above the critical point of the model. The state in all cases was found to be in the paramagnetic phase, as was found in Ezaki et al. (2020) [9].

Since Ising temperature was expected to correlate with entropy and disorder in the models, and because of the relevance of the notion of complexity in the analysis of brain data, we studied two proxies (upper bounds) of algorithmic complexity, one closely related to Shannon entropy (LZW), the other more directly connected with the concept of Kolmogorov complexity (BDM). We did this first, starting from the data itself and then also working with synthetic data generated from the personalized Ising models. Complexity metrics derived from the data related only weakly (but significantly at the group level) with temperature and condition, presumably due to the limited amount of data per participant and condition, and we cannot discount the possibility of paradoxical effects of LSD in some subjects leading to decreases of complexity. However, it is well known that measures that estimate complexity are sensitive to data length [67, 93], and this is a limitation in this study because BOLD data is intrinsically slowly varying and the time series are not very long. To compensate for this, we explored model-derived LZW and BDM complexity estimates using synthetic data. We found these to be monotonic functions of temperature (Figure 3 and 5) that correlated strongly with the condition.

Figure 3 (bottom right) displays the monotonic behavior of BDM complexity with respect to temperature, with a very sharp increase and then almost a plateau. This suggests the bound on complexity magnitude, whereas the slope of increase should be correlated with model parameters describing the simulation condition. One limitation in the BDM calculation is that, as the size of matrices increases, BDM gets gradually closer and closer to an entropy measurement as we use precomputed CTM for smaller matrices. Moreover, it is important to note that, in our Ising and complexity analysis, we are bounded to binary data, which makes the analysis contingent on how meaningful the binarized version of the data is.

Overall, our results are consistent with the notion that psychedelics drive brain dynamics into a more disordered state and, in our modeling framework, away from criticality. The RElaxed Beliefs Under pSychedelics (REBUS) framework [1]) addresses the phenomenology and physiology of psychedelic states by integrating mechanistic insights with the free-energy principle (FEP, [94, 95]). The FEP is a formulation based on causal statistical theory that accounts for action, perception, and learning, and the entropic brain hypothesis (EBH [32]), which associates entropy of spontaneous brain activity with the richness (vividness and diversity) of subjective experience (and posits that psychedelics increase both).

### Relation to REBUS model and criticality

The mechanisms of action of psychedelics are believed to begin with stimulation of a specific serotonin receptor subtype (5-HT_2A_) on cortical layer V pyramidal neurons, i.e., this is the control signal that results in increased neuronal excitability and dysregulated cortical population-level activity [96]. Through this action, psychedelics are believed to disrupt the regular functioning of various high-level system properties—e.g., major cortical oscillatory rhythms and the integrity of large-scale networks—that are hypothesized to encode the precision-weighting of internal models—i.e., priors, beliefs, or assumptions. According to the so-called ‘REBUS’ model [1], psychedelics disrupt models sitting atop the modeling hierarchy, with direct consequences on experience. Interference with microcircuitry associated with high-level models may be expected to release lower-level models that would otherwise be suppressed.

In [3], we hypothesized that increased apparent (Shannon) complexity (entropy) of spontaneous activity is a logical corollary of this action, as apparently complex, bottom-up uncompressed data more strongly dominates brain activity. Such increases have indeed been observed using measures such as LZ [97] and are also seen in our data. The present study extends on previous work, however, by incorporating algorithmically-oriented methods such as BDM. Our results agree with earlier work pointing to the usefulness of complexity metrics and the hypothesis that signal complexity should increase with increased system temperature, and that psychedelics should increase both [2]. With regard to the criticality perspective, in Ruffini & Lopez-Sola (2022), we proposed that a modeling system tracking structured data in the world (a wakeful brain) will display dimensionality reduction and criticality features that can be used empirically to quantify the structure of the program run by an agent [3]. Criticality features are seen to result from the collapse of a dynamical system (the dynamical brain) into a low dimensional space associated with identified regularities in complex data—the hallmark of an algorithmic compressive system. If tracking world data using models keeps the modeling dynamics on a reduced, lower-dimensional manifold, psychedelics may be said to “lift” up the enslaved dynamics into higher dimensions. Our finding of the resting brain state being above criticality and driven further away from the critical point under the influence of LSD is consistent with this view.

As discussed in the context of the algorithmic information theory of consciousness (KT) [2, 3], psychedelics will shift the dynamics of an agent system tracking world data to a less constrained state further away from criticality, and hence produce more complex signals. The above is in line with the idea that the brain operates near a critical point (v. [24, 98, 99] and references therein), and that psychedelics move dynamics into a more disordered state [1]. However, at least in our model and as already described in previous work using similar methods [8, 9], we found that the resting brain in the placebo condition was already above the critical point—that is, the resting, the wakeful brain is in a supercritical state as observed through the Ising framework lens. This is consistent with the idea that, while mutual information peaks at the critical temperature, information flow in such systems peaks in the disordered (paramagnetic) phase [100], which is in apparent contradiction with other studies suggesting a subcritical nature of healthy brain dynamics [1]. We note here in passing that the term “criticality” is used differently by different authors, e.g., dynamical vs. statistical criticality, whose relation is the subject of current research [99]. At least with the data in this study, the statistical criticality metrics developed here—the supercritical system temperature of an Ising model—, provided the best correlate of experimental condition.

Finally, with regard to the relation of these metrics with subjective experience as measured by questionnaires, some weak correlations were found at the trend level with all metrics (temperature and complexity metrics), presumably because of the limited number of data and the inherent heterogeneity of self-reports and that these particular ratings were done post-hoc without precise reference to the scanning periods. Specifically designed neurophenomenological methods probing the structure of experience during altered states of consciousness may provide better correlates in future studies [3].

## Conclusions

In this paper, we have studied criticality features of brain dynamics under LSD in a within-subject study using the Ising model formalism and algorithmic complexity using Lempel-Ziv and the Block Decomposition methods. Personalized Ising models were created using BOLD fMRI data by adjusting the model temperature for each subject and condition in relation to the group average model. We found that the effects of LSD translate into increased BOLD signal complexity and Ising temperature, in agreement with earlier findings and predictions from existing theories of the effects of psychedelics such as the relaxed beliefs under psychedelics (REBUS), the anarchic brain hypothesis [1], and the algorithmic information theory of consciousness (KT) [2, 3]. More precisely, in agreement with earlier work [9], we found that the system in the placebo condition was already in the paramagnetic phase—above the critical point—, with ingestion of LSD resulting in a shift away from criticality into a more disordered state. These results must be interpreted with care, as the Ising framework used here provides a simple one-parameter model for fitting the individual data. Other studies, using different methods, suggest that LSD shifts system dynamics closer to criticality [101, 102], while, for example, using similar methods to ours, higher fluid intelligence has been associated with closer to critical resting-state neural dynamics [9]. This highlights the question of whether modeling choices such as what data and data features to model (e.g., BOLD FC or electrophysiological data), modeling scale (micro, meso or macroscale), how to model it (e.g., Ising, Hopf, or neural mass models) can impact conclusions on the critical features of the system. Further work is needed to reconcile these results, which may stem from both these modeling choices and associated criticality metrics [103]. The Ising formalism provides us with a parsimonious method to infer structural connectivity from functional data. The Ising connectome, we found, is related to the structural (dMRI) connectome but differs in two important ways. First, it includes stronger homotopic connectivity, which, as we saw, is weakened during LSD. Second, it is a signed connectome, and the negative connections play an important dynamical role.

## Acknowledgments

GR, GiD, DL-S, ND have received funding from the European Research Council (ERC Synergy Galvani) under the European Union’s Horizon 2020 research and innovation programme (grant agreement No 855109) and under European Union’s Horizon 2020 research and innovation programme under grant agreement No 101017716 (Neurotwin). RLC-H is supported by the Ralph Metzner Distinguished Professorship endowment. GuD is supported by the ERC Advanced Grant DYSTRUCTURE (295129), the Spanish Research Project PSI2016-75688-P, the European Union’s Horizon 2020 Framework Programme for Research and Innovation under the Specific Grant Agreement No. 785907 (Human Brain Project SGA2), the European Union’s Horizon 2020 research and innovation programme (grant agreement No 855109), and under European Union’s Horizon 2020 research and innovation programme under grant agreement No 101017716 (Neurotwin). NAK has received funding from the European Union under the Horizon 2020 programme (MultipleMS grant agreement 733161 and DECISION Grant agreement 847949). FR is supported by the Ad Astra Chandaria foundation. AP-A was supported by the Human Brain Project SGA3 (945539).

## Data and code availability

Code for Ising modeling can be found at https://github.com/giulioruffini/LSD-paper-code-Ruffini-et-al.-2022, and for LZW analysis at https://github.com/giulioruffini/StarLZW. The code for fMRI analysis is available at https://github.com/decolab/cb-neuromod. The multimodal neuroimaging data from the experiment is available upon request from (contact Dr. Robin Carhart-Harris). BDM calculation programs in different languages are available on the algorithm dynamics lab website, software page, https://www.algorithmicdynamics.net/software.html.

## Contributions

GR: conceptualization, methodology, software, writing the original manuscript; GiD: data processing, software, manuscript contributions; DLS: statistical analysis support; ND: conceptualization, software, data processing; FER: manuscript revision, conceptualization; NAK: data analysis (BDM), conceptualization, manuscript revision; APA: conceptualization; MLK: data processing; RC-H: experimental conceptualization, data collection, manuscript revision; GuD: conceptualization, manuscript revision.

## Supporting Information

### A List of AAL areas

Here we report the list of AAL areas in the specific order they were used in this study (Table SI.1).

**Table SI.1.** Regions of interest from AAL atlas. The AAL areas were ordered in the following way: first all the areas in the left hemisphere (LH), and then all the areas in the right hemisphere (RH) in reverse order, so that they are “symmetrized” with respect to the contralateral hemisphere.

| idx | ROI name |  | idx | ROI name |  |
| --- | --- | --- | --- | --- | --- |
| 1 | Precentral | LH | 46 | Temporal_Inf | RH |
| 2 | Frontal_Sup | LH | 47 | Temp_Pole_Mid | RH |
| 3 | Frontal_Sup_Orb | LH | 48 | Temporal_Mid | RH |
| 4 | Frontal_Mid | LH | 49 | Temp_Pole_Sup | RH |
| 5 | Frontal_Mid_Orb | LH | 50 | Temporal_Sup | RH |
| 6 | Frontal_Inf_Oper | LH | 51 | Heschl | RH |
| 7 | Frontal_Inf_Tri | LH | 52 | Thalamus | RH |
| 8 | Frontal_Inf_Orb | LH | 53 | Pallidum | RH |
| 9 | Rolandic_Oper | LH | 54 | Putamen | RH |
| 10 | Supp_Motor | LH | 55 | Caudate | RH |
| 11 | Olfactory | LH | 56 | Paracentral_Lob | RH |
| 12 | Frontal_Sup_Med | LH | 57 | Precuneus | RH |
| 13 | Frontal_Med_Orb | LH | 58 | Angular | RH |
| 14 | Rectus | LH | 59 | SupraMarginal | RH |
| 15 | Insula | LH | 60 | Parietal_Inf | RH |
| 16 | Cingulum_Ant | LH | 61 | Parietal_Sup | RH |
| 17 | Cingulum_Mid | LH | 62 | Postcentral | RH |
| 18 | Cingulum_Post | LH | 63 | Fusiform | RH |
| 19 | Hippocampus | LH | 64 | Occipital_Inf | RH |
| 20 | ParaHippo | LH | 65 | Occipital_Mid | RH |
| 21 | Amygdala | LH | 66 | Occipital_Sup | RH |
| 22 | Calcarine | LH | 67 | Lingual | RH |
| 23 | Cuneus | LH | 68 | Cuneus | RH |
| 24 | Lingual | LH | 69 | Calcarine | RH |
| 25 | Occipital_Sup | LH | 70 | Amygdala | RH |
| 26 | Occipital_Mid | LH | 71 | ParaHippo | RH |
| 27 | Occipital_Inf | LH | 72 | Hippocampus | RH |
| 28 | Fusiform | LH | 73 | Cingulum_Post | RH |
| 29 | Postcentral | LH | 74 | Cingulum_Mid | RH |
| 30 | Parietal_Sup | LH | 75 | Cingulum_Ant | RH |
| 31 | Parietal_Inf | LH | 76 | Insula | RH |
| 32 | SupraMarginal | LH | 77 | Rectus | RH |
| 33 | Angular | LH | 78 | Frontal_Med_Orb | RH |
| 34 | Precuneus | LH | 79 | Frontal_Sup_Med | RH |
| 35 | Paracentral_Lob | LH | 80 | Olfactory | RH |
| 36 | Caudate | LH | 81 | Supp_Motor | RH |
| 37 | Putamen | LH | 82 | Rolandic_Oper | RH |
| 38 | Pallidum | LH | 83 | Frontal_Inf_Orb | RH |
| 39 | Thalamus | LH | 84 | Frontal_Inf_Tri | RH |
| 40 | Heschl | LH | 85 | Frontal_Inf_Oper | RH |
| 41 | Temporal_Sup | LH | 86 | Frontal_Mid_Orb | RH |
| 42 | Temp_Pole_Sup | LH | 87 | Frontal_Mid | RH |
| 43 | Temporal_Mid | LH | 88 | Frontal_Sup_Orb | RH |
| 44 | Temp_Pole_Mid | LH | 89 | Frontal_Sup | RH |
| 45 | Temporal_Inf | LH | 90 | Precentral | RH |

### B Changes in connectivity

**Fig SI.2.**
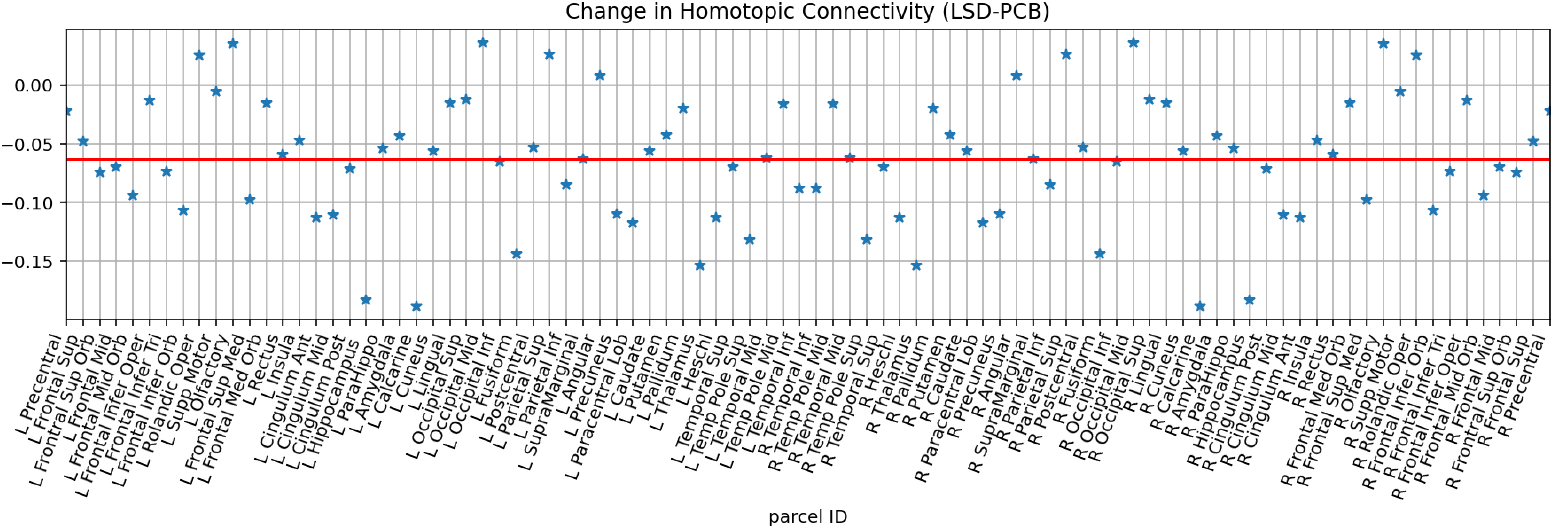
Changes in homotopic connectivity in Ising connectivity matrix (LSD-placebo). The largest decreases can be observed in the Calcarine (ID 19), Hippocampus (22), Thalamus (39), and Fusiform (28) areas. The largest increases are found in the Rolandic Operculum (9), Olfactory (11), Occipital Mid (26), and Parietal Sup (30) areas. The red line indicates the mean across all points (negative in this case, since there is an average decrease in homotopic connectivity).

### C Susceptibility per parcel

**Fig SI.3.**
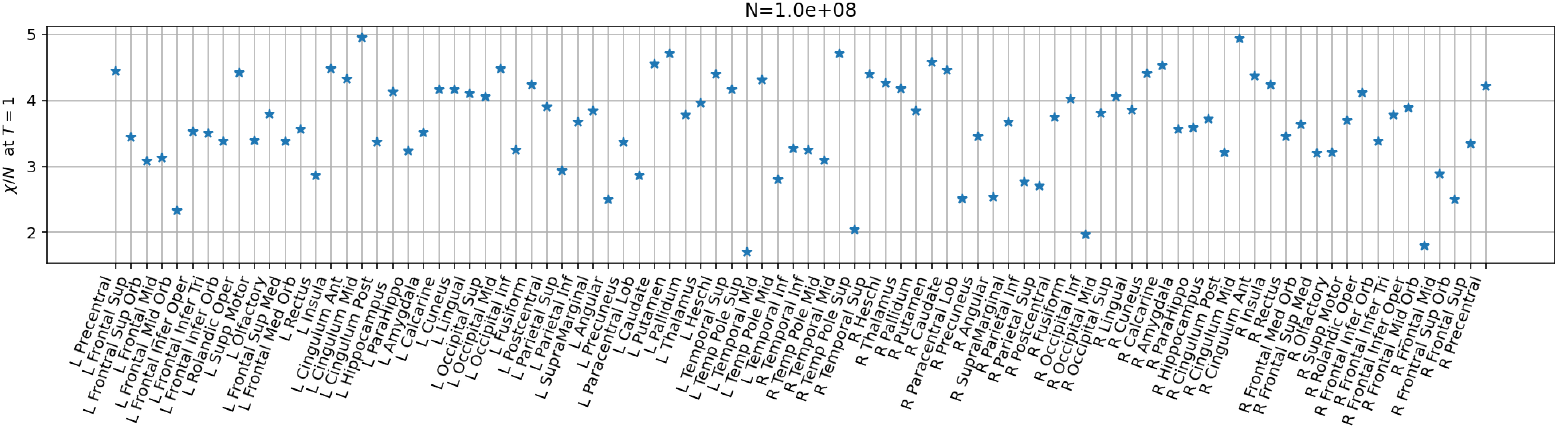
Susceptibility per parcel at *T* = 1. For each parcel, the susceptibility of each parcel at the nominal *T* = 1 temperature of the global archetype Ising model is provided.

**Fig SI.4.**
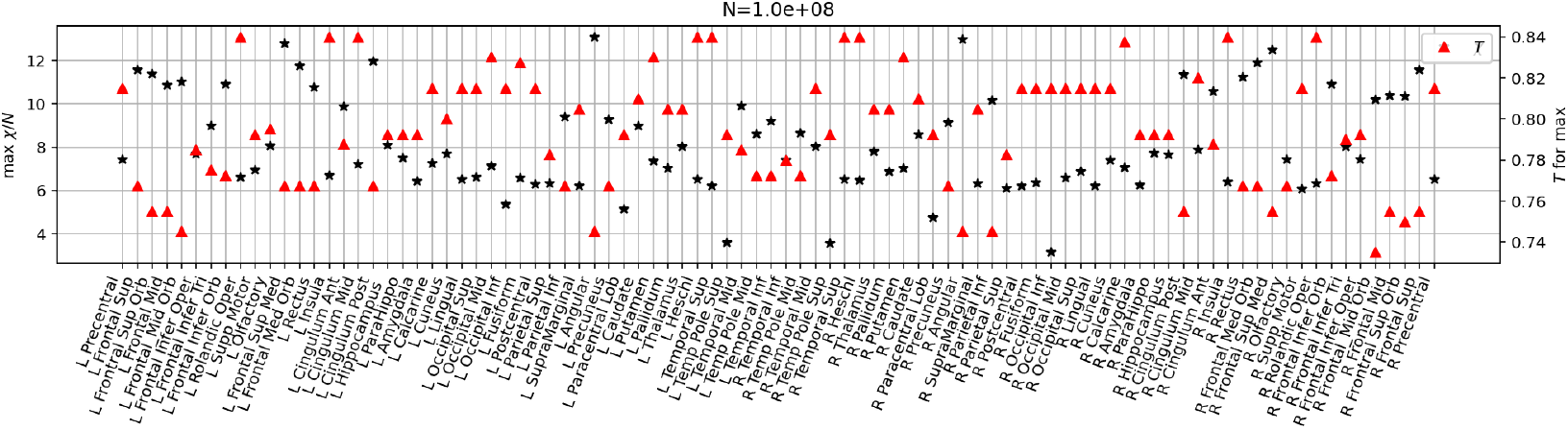
Maximum susceptibility per parcel and corresponding temperature. For each parcel, the maximum susceptibility (black asterisk) and the temperature at which it is attained (red triangle) are provided for the global Ising archetype model.

### D Empirical FC, all data and delta

**Fig SI.5.**
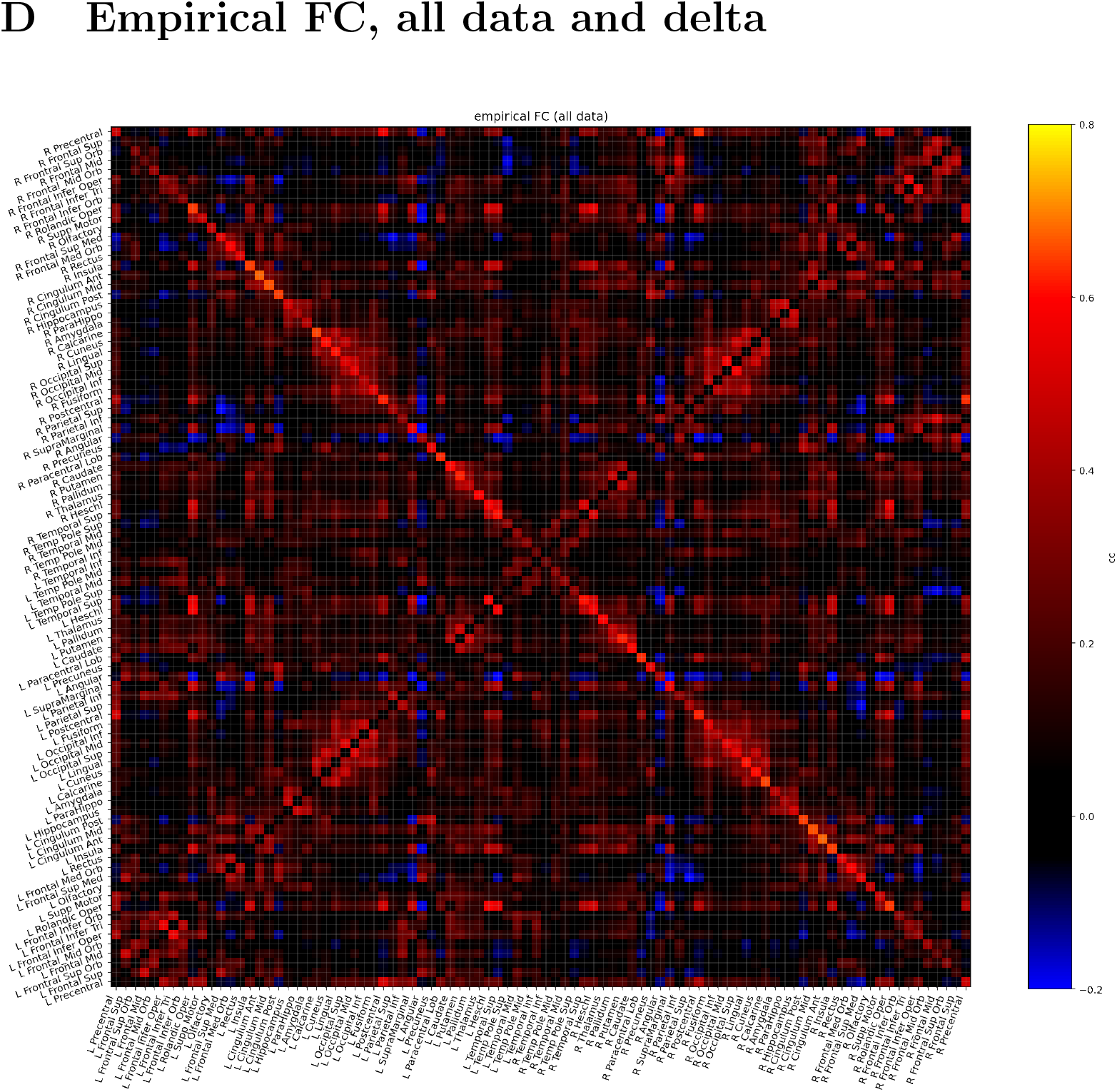
Empirical FC, all data. Pearson correlation between the binarized average activity of each parcel.

**Fig SI.6.**
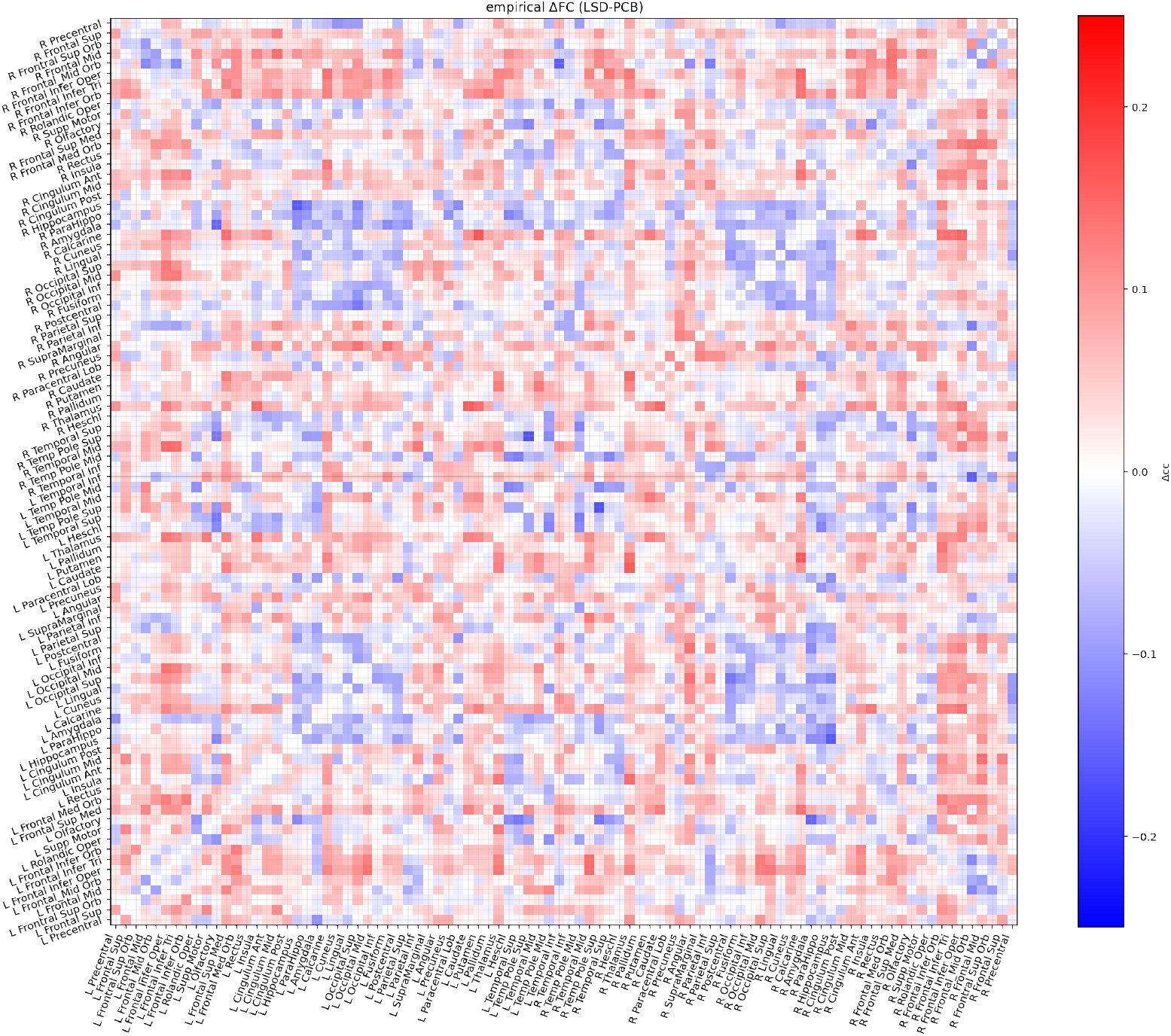
Change in empirical FC between two conditions (LSD minus placebo).

### E Ising and dMRI Connectomes

**Fig SI.7.**
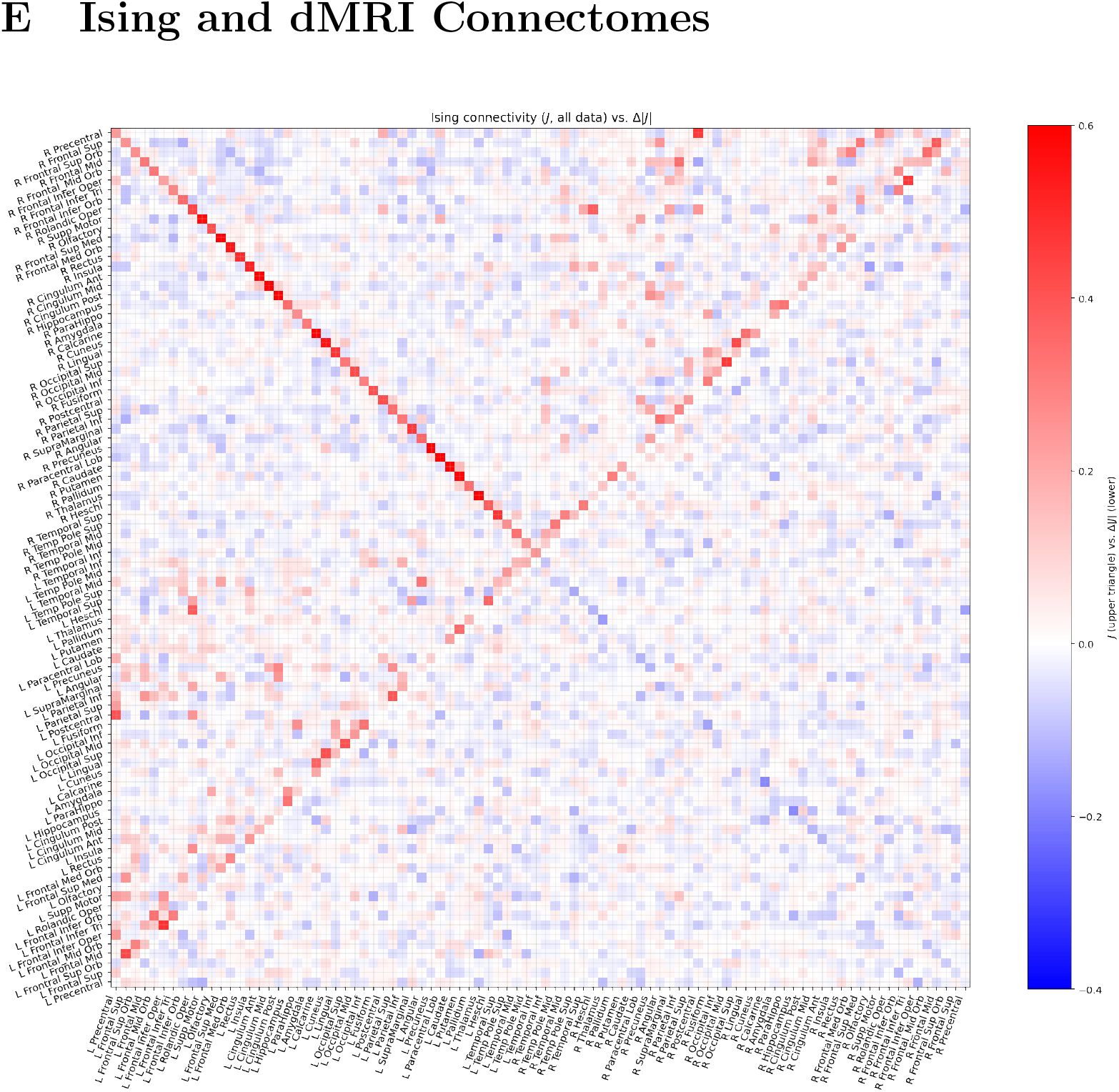
**Left:** Global archetype (combining all data) Ising connectivity *J* (upper left triangle), and the change in link strength (Δ|*J*|, LSD-placebo) (lower right triangle). The down-trending diagonal values reflect strong inter-hemispheric homotopic connectivity and its change (decrease in LSD condition). **Top right:** difference in connectivity magnitude between the placebo and LSD archetypes, with a visible loss in homotopic connectivity. By construction, the connectomes have zeros in the diagonal elements (e.g., *J_ii_* = 0).

**Fig SI.8.**
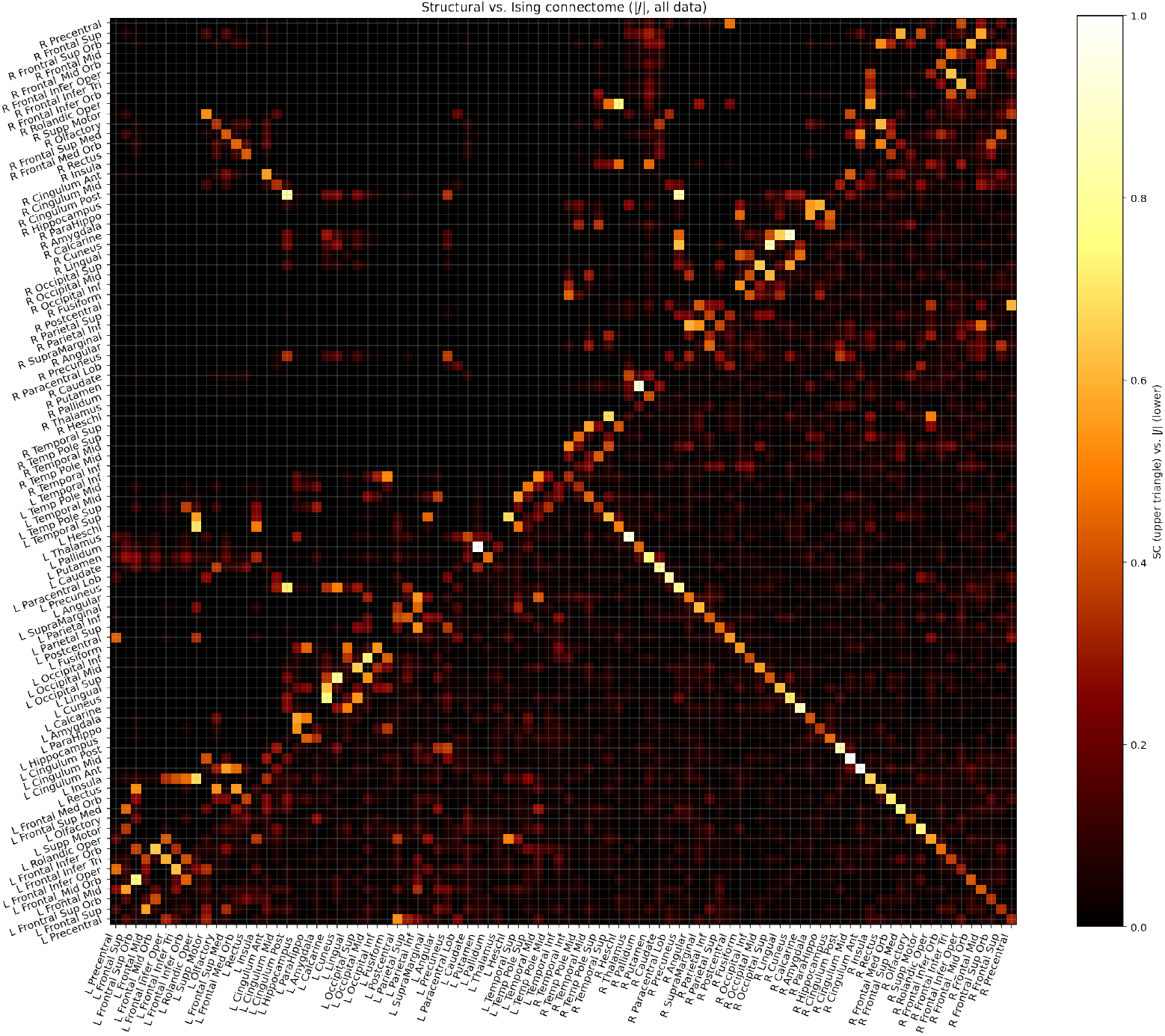
Ising-derived connectome: AAL structural connectome obtained from dMRI (from [85]) (upper left triangle) and unsigned global archetype Ising connectivity (lower right triangle).

### F Network plots

Here we provide plots of the Ising connectivity magnitude change (LSD-PCBO).

**Fig SI.9.**
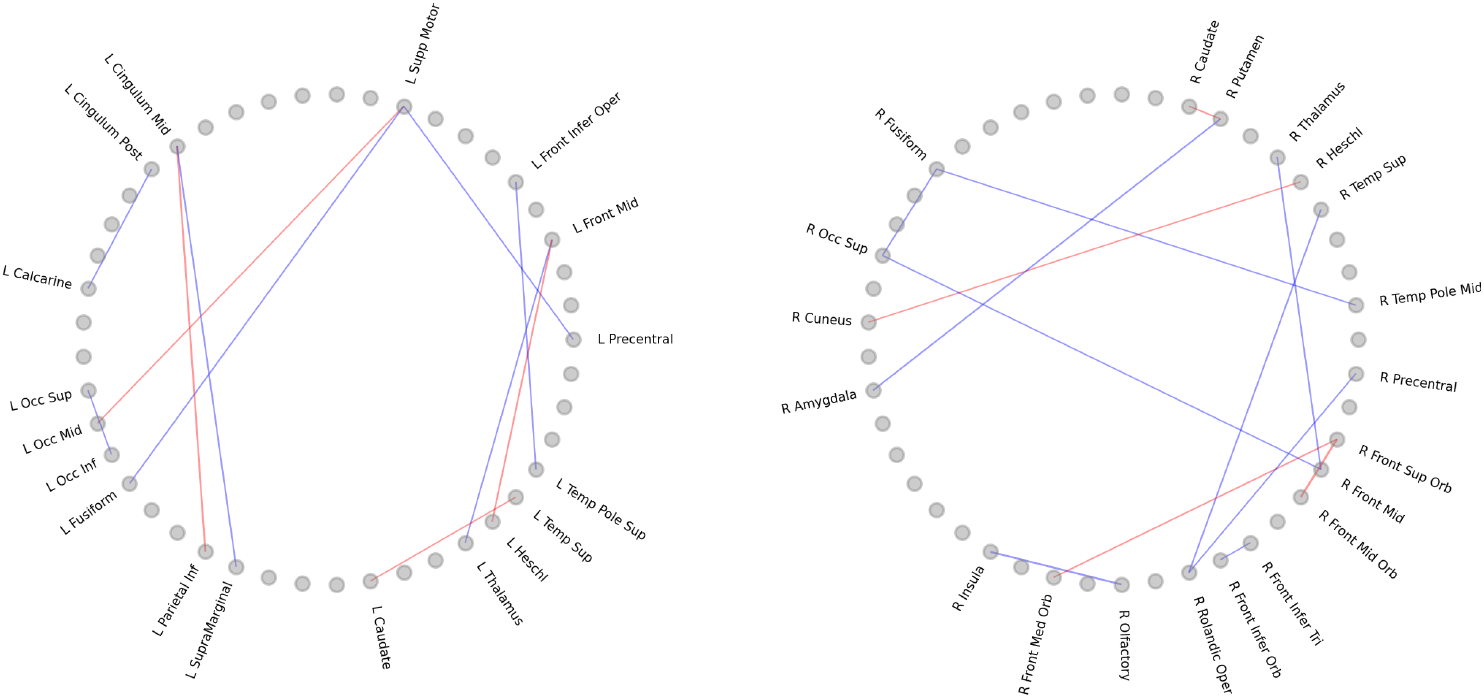
Intra-hemispheric network plots highlighting changes (Δ|*J*|, LSD-PCBO) higher than 0.11 in the left (on the left) and right (right) hemispheres. Red (blue) = positive (negative) connections.

**Fig SI.10.**
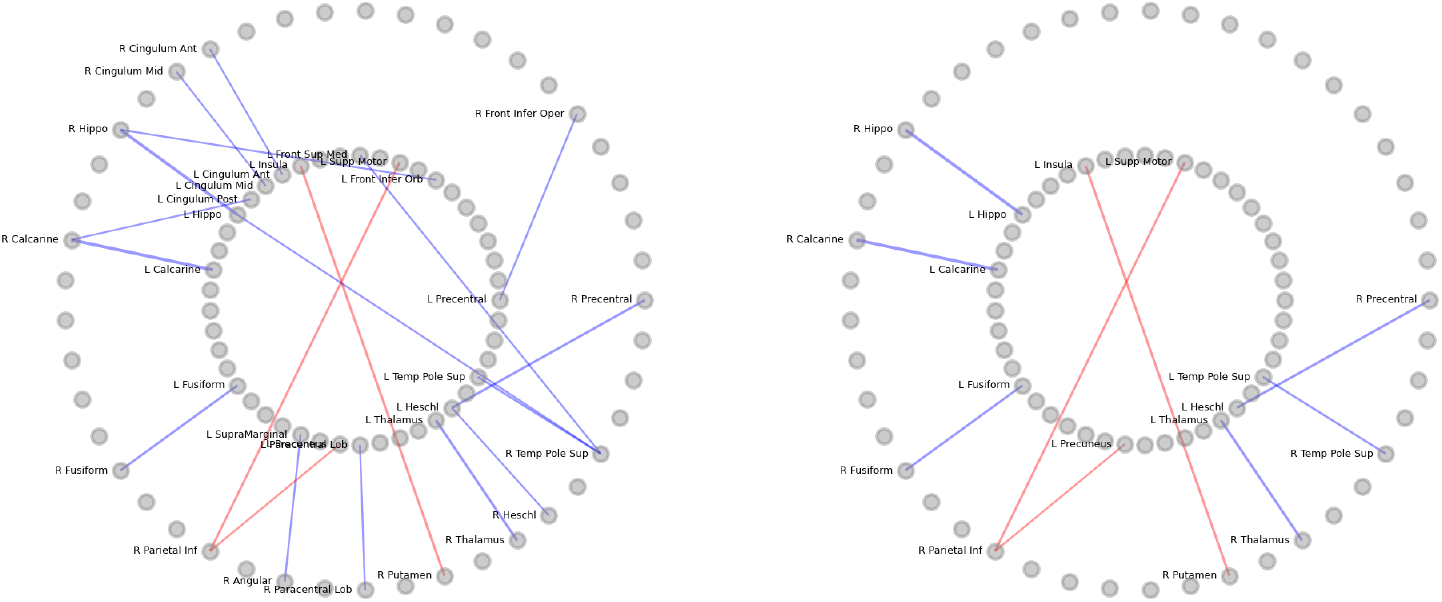
Inter-hemispheric network plots highlighting changes (Δ| *J*|, LSD-PCBO) higher than 0.11 (left) and 0.12 (right). The inner (outer) circle displays the left (right) hemisphere nodes. Red (blue) = positive (negative) connections.

### G Group FC for each condition (empirical vs. Ising)

**Fig SI.11.**
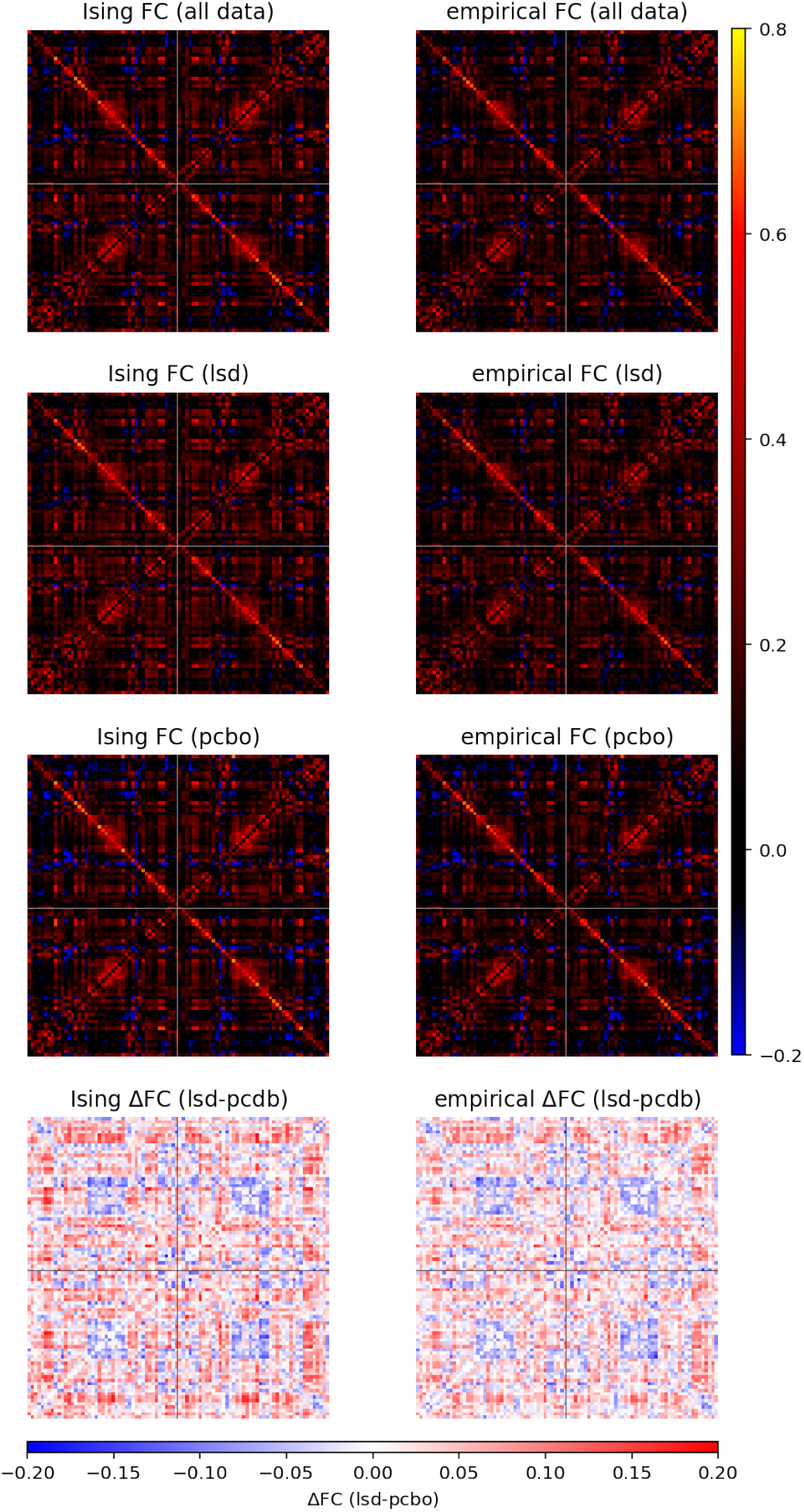
FC for each condition, empirical vs. Ising generated (LSD, placebo and difference). **Left column:** FC generated by each archetype (LSD and placebo) and difference on the bottom. We note that 1 bit quantization produces a loss of correlation as compared to unbinarized correlation. **Right column:** Empirical FC for each condition and difference (bottom) from binarized data. The Pearson correlation between the empirical and Ising FC (all data) is 0.99, and between the difference (delta FC of LSD-placebo, empirical vs. Ising) is 0.96.

### H Homotopic FC and FC negativity. FC for each subject and condition

Figure SI.12 provides boxplots of the summed homotopic FC (FC of a subset of homotopic links) and the negativity of each FC matrix. We define negativity simply by the percent weight of negative weights,

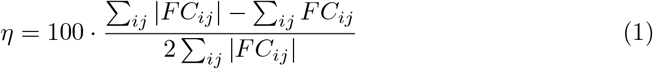

**Fig SI.12.**
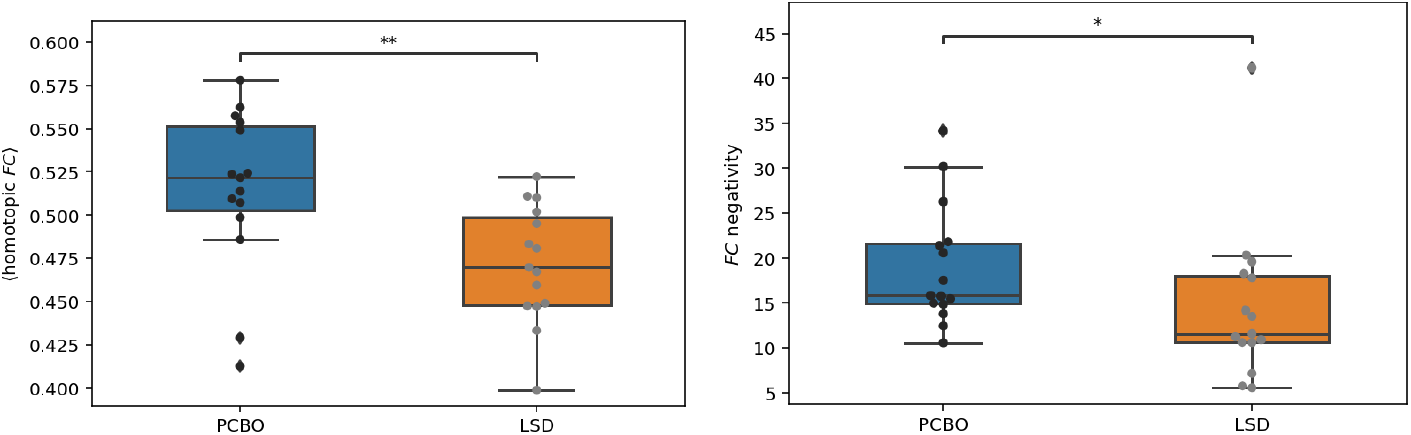
Homotopic FC for the two groups and percent of weight of negative values in FC (negativity). Left: Homotopic connectivity (paired Wilcoxon statistic=10.0, p=0.003); Right: FC negativity (paired Wilcoxon statistic=16.0, p=0.010)

In the next figures, we provide the data for each subject (Fig SI.13).

**Fig SI.13.**
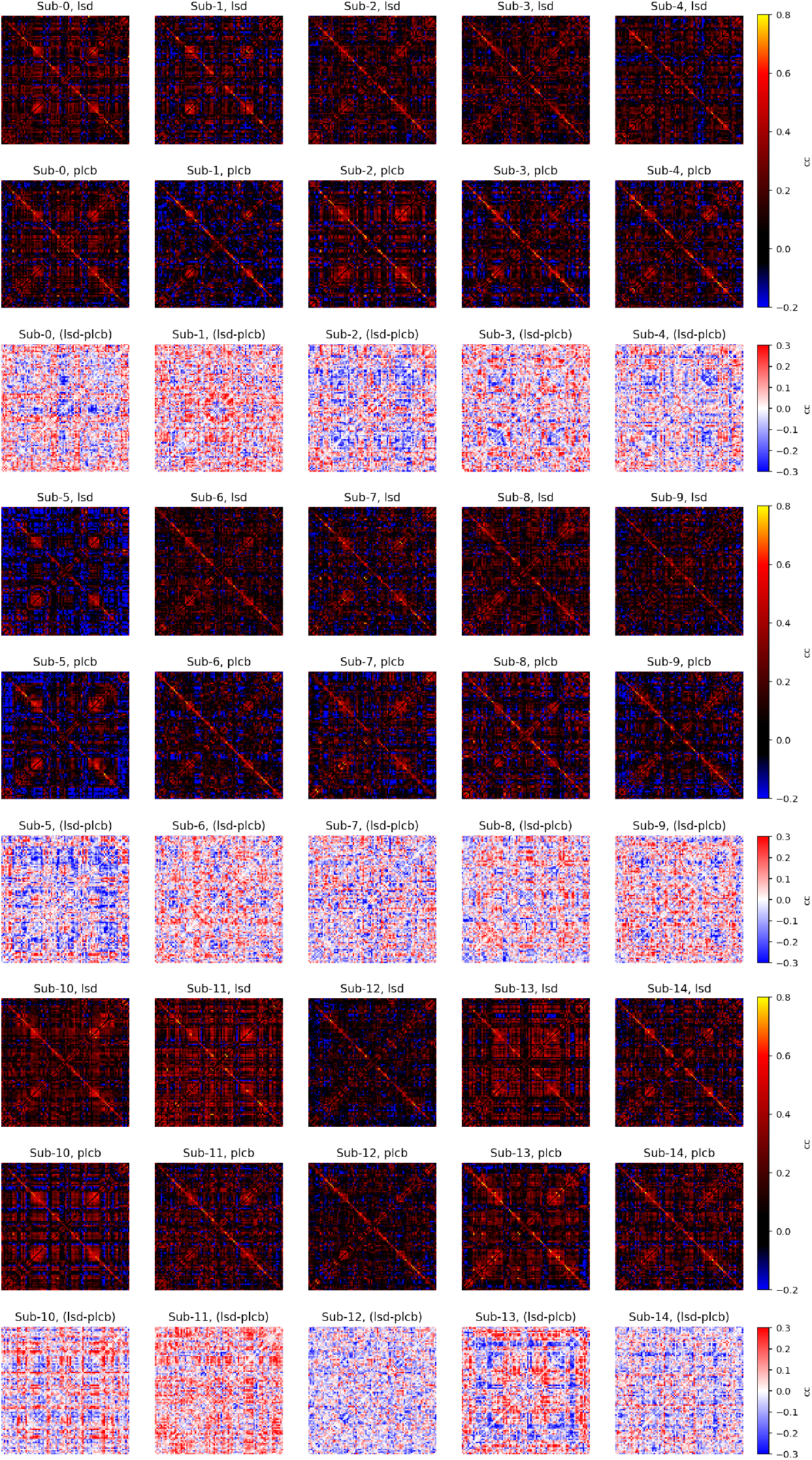
FC of binarized empirical data for each subject. LSD (top), Placebo (middle), and the difference between the two (bottom). The color scale is adjusted to display weaker but numerous negative links in FC.

### I Homotopic and link susceptibility

Figure SI.14 provides plots of susceptibility for the placebo data archetype and the effect of reducing homotopic connectivity by 20%, which decreases the critical temperature and leads to loss of FC.

**Fig SI.14.**
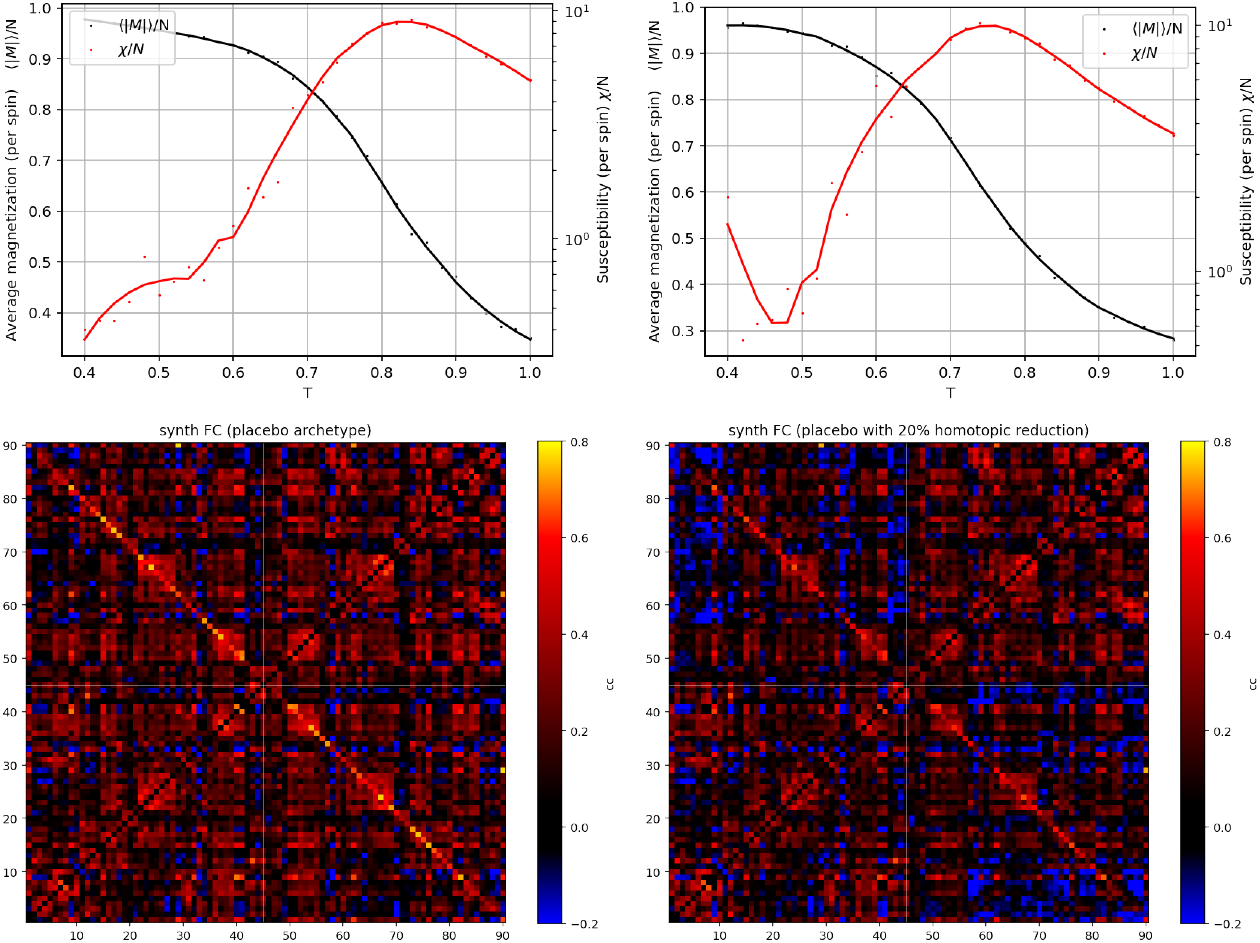
Impact of reduced homotopic links strength: Top left, susceptibility and magnetization plot of placebo archetype (*T*_c_ = 0.84); top right: susceptibility and magnetization plot after reducing homotopic link strength by 20% (*T*_c_ = 0.74). Bottom: respective FC of synthetic data generated by each model at *T* = 1.

Next, we analyze the concept of link susceptibility. The link susceptibility

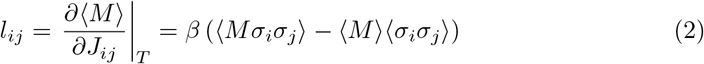

provides information about the sensitivity to additive changes to a particular link in the system. To understand the impact of changes in the scale of a particular link (*J* → *J* + *αJ*), we use the scaled quantity *L_ij_* = *l_ij_J_ij_*, which provides a metric on the susceptibility of the mean magnetization of the model to multiplicative changes in archetype link strength,

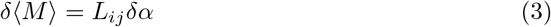

Figure SI.15 provides the link and scaled link susceptibility for the placebo archetype near the critical temperature.

**Fig SI.15.**
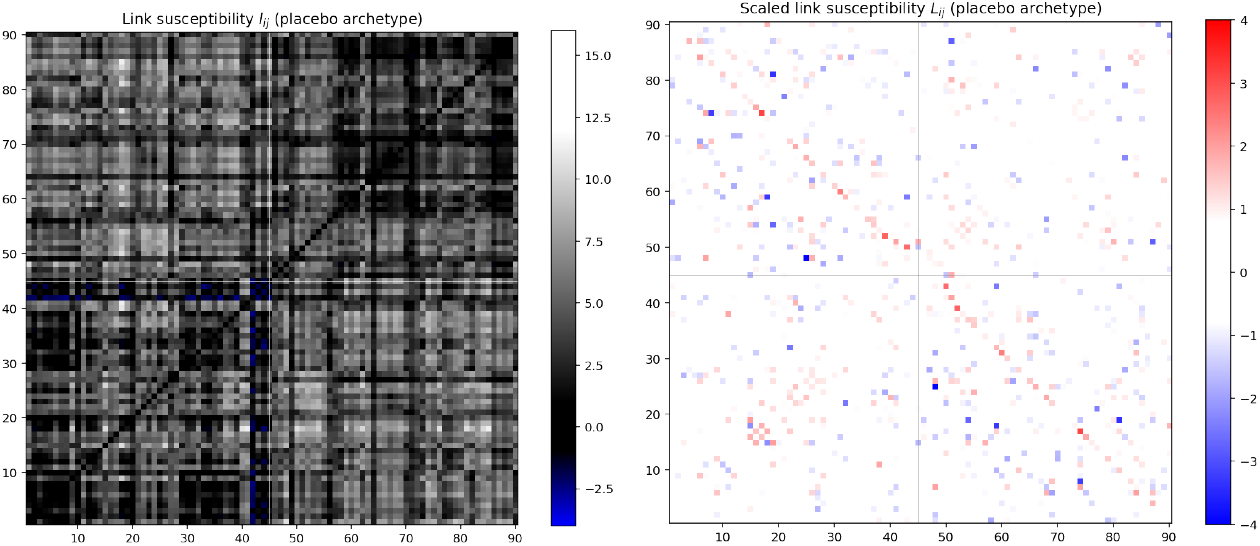
Link susceptibility: Left, link susceptibility plot of placebo archetype; right, scaled link susceptibility.

### J Correlation between metrics

We checked the correlation between features extracted from data as shown in Figure SI.16.

**Fig SI.16.**
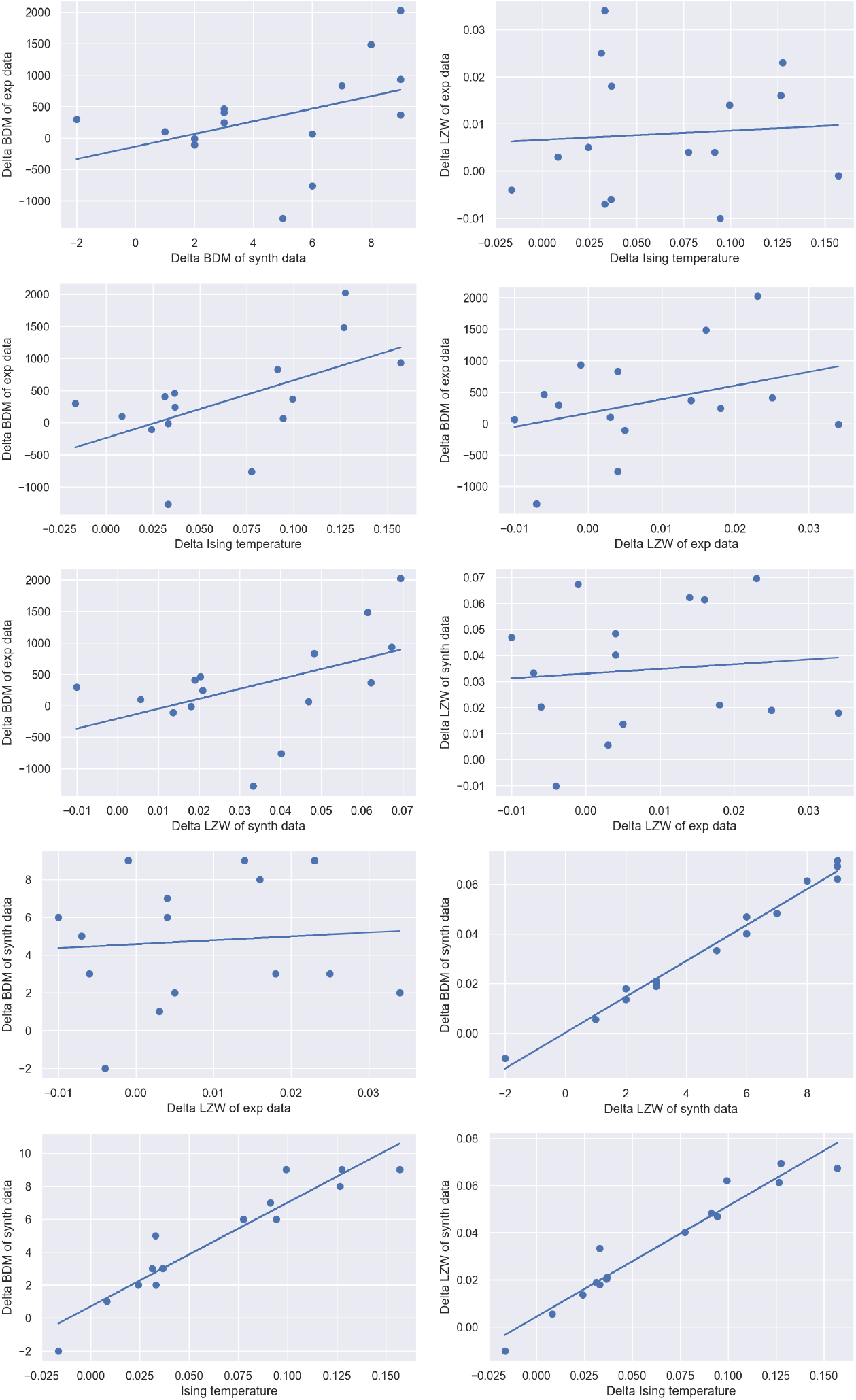
Correlation scatter plots between the metrics. Correlation plots between the metrics, i.e., delta LZW complexity of the simulated and empirical data, delta Ising temperature, and delta BDM complexity of the simulated and empirical data.

### K Subjects questionnaire scores and their correlation with metrics

**Table SI.16.**
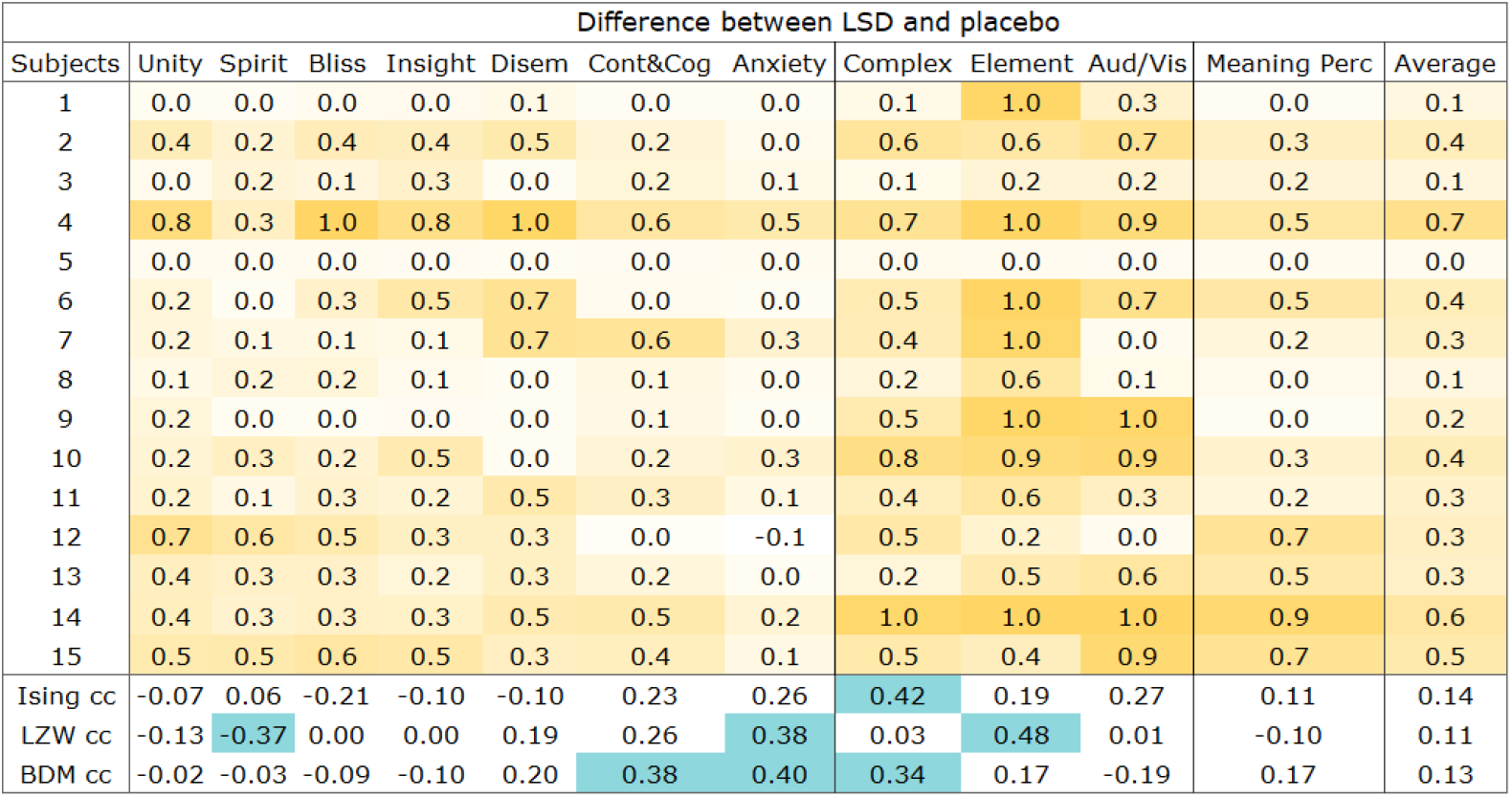
ASC questionnaire scores and their correlation with features extracted from data. The higher scores are colored in darker yellow, whereas Pearson correlation coefficients higher in absolute value than 0.30 are colored in blue. Full names of ASC questionnaire categories: Experience of unity, Spiritual experience, Blissful state, Insightfulness, Disembodiment, Impaired control and cognition, Anxiety, Complex imagery, Elementary imagery, Audio-visual synesthesiae, Changed meaning of percepts.

**Table SI.16.** VAS questionnaire scores and their correlation with features extracted from data. The higher scores are colored in darker yellow, whereas Pearson correlation coefficients higher in absolute value than 0.30 are colored in blue.

| Average difference between LSD and placebo for session 1 and 2 |  |  |  |  |  |  |  |
| --- | --- | --- | --- | --- | --- | --- | --- |
| Subject | Intensity | Simple | Complex | Emotional Arousal | Positive Mood | Ego Dissolution | Average |
| 1 | 11 | 16 | 11 | 3 | 10 | 0 | 8 |
| 2 | 12 | 10 | 7 | 12 | 4 | 16 | 10 |
| 3 | 11 | 5 | 1 | 3 | 2 | 0 | 3 |
| 4 | 13 | 20 | 20 | 11 | 10 | 2 | 13 |
| 5 | 9 | 0 | 1 | 12 | -1 | 0 | 4 |
| 6 | 15 | 17 | 17 | 6 | 10 | 6 | 12 |
| 7 | 11 | 1 | 1 | 12 | 0 | 1 | 4 |
| 8 | 9 | 6 | 3 | 10 | 1 | 0 | 5 |
| 9 | 4 | 4 | 0 | 5 | 7 | 0 | 3 |
| 10 | 13 | 11 | 3 | 10 | 0 | 0 | 6 |
| 11 | 11 | 9 | 10 | 10 | 12 | 3 | 9 |
| 12 | 19 | 15 | 12 | 19 | 19 | 20 | 17 |
| 13 | 15 | 10 | 4 | 15 | 5 | 11 | 10 |
| 14 | 19 | 19 | 19 | 18 | 17 | 17 | 18 |
| 15 | 15 | 5 | 6 | 14 | 8 | 11 | 10 |
| Ising cc | 0.10 | -0.15 | -0.26 | 0.31 | -0.33 | 0.11 | -0.07 |
| LZW cc | -0.07 | 0.34 | 0.35 | -0.38 | 0.06 | -0.22 | 0.05 |
| BDM cc | 0.31 | 0.15 | 0.19 | 0.39 | 0.02 | 0.11 | 0.22 |

